# Effects of forest cover, temperature and flower abundance on cashew pollinators in the northern Western Ghats, India

**DOI:** 10.1101/2025.11.11.686480

**Authors:** Aditya Satish, Vishal Sadekar, Nandita Madhu, Sanath Manimoole, H Rama Narayanan, Anushka Rege, Rohit Naniwadekar

## Abstract

Global declines in pollinators pose serious risks for biodiversity, ecosystem functioning, and human well-being. Understanding how landscape structure and climate shape pollinator communities is critical, yet studies that jointly assess these factors in human-dominated tropical landscapes remain scarce. Here, we investigate how forest cover, distance to forest patches, flower abundance, and temperature influence pollinator communities in monoculture cashew plantations that are rapidly expanding in the northern Western Ghats, part of the Western Ghats-Sri Lanka global biodiversity hotspot. We conducted 210 half-hour observation sessions across 30 cashew orchards in Dodamarg, India. At the community level, flower abundance and forest cover positively influenced pollinator richness, number of pollinators, and number of visits. The number of visits showed a unimodal response to temperature, with peaks occurring within narrow, taxon-specific ranges, lower temperatures for birds and higher for insects, likely reflecting physiological constraints. At the species level, flower abundance and temperature emerged as key predictors of occurrence, while three butterfly species occurred more frequently near forest patches. Contrary to expectations, proximity to forest did not reduce ambient orchard temperatures, suggesting limited forest-mediated thermal buffering in our site. Together, these findings highlight the vulnerability of pollination services to both land-use change and rising temperatures. Ongoing conversion of natural forests to cashew monocultures, coupled with climate warming, may erode pollinator communities and jeopardize crop yields in tropical agroecosystems. More broadly, our results underscore the importance of conserving remnant forests and accounting for thermal sensitivity of pollinators to sustain ecosystem services in changing landscapes worldwide.

## 1. Introduction

Pollination is an essential ecosystem service that supports both ecosystem functioning and agricultural production. Pollinators are critical for sexual reproduction in nearly 90% of angiosperms (Ollerton et al., 2011), thereby contributing directly to fruit set in both wild and cultivated plants (Garibaldi et al., 2011, 2013). Humans rely heavily on crop-pollination services, with their global economic value estimated at up to 387 billion USD annually (Porto et al., 2020). However, these services are increasingly threatened by large-scale declines in pollinator diversity (Kearns et al., 1998; Steffan-Dewenter et al., 2002, 2005). These declines have largely been driven by land-use change and climate change (Potts et al., 2010), which alter (a) pollinator community composition and (b) the spatial distribution and temporal activity of pollinators, thereby affecting their interactions with flowering plants (Rafferty, 2017). Identifying factors that sustain pollinator diversity and abundance is crucial for both biodiversity conservation and human well-being.

Approximately 30% of global forest cover has been lost to human-driven deforestation over the last three centuries, primarily due to the growing demand for agricultural land (Ritchie, 2021). The hypotheses proposed to explain the negative impacts of land-use change on pollinators are linked to the habitat, feeding, and breeding biology of pollinator species. Since many pollinator species depend on forests for nesting, refuge, and food, the loss of forests can deprive pollinators of critical food and nesting resources (Ulyshen et al., 2023). Compared to agricultural lands, forests may provide a higher diversity of floral sources, which can support greater pollinator diversity in the classic ‘diversity-begets-diversity’ fashion (Jha & Vandermeer, 2010; Whittaker, 1972). Beyond floral resources, pollinators also use non-floral resources at different stages of their life cycle—for example, lepidoteran larvae feed on leaves of forest plants, while many bees nest in tree cavities (Ulyshen et al., 2023). These resources tend to be more abundant in undisturbed, contiguous forests. Consistent with this expectation, studies show positive association between pollinator diversity and (a) forest patch size (Aguirre & Dirzo, 2008; Smith & Mayfield, 2018) and (b) the availability of deadwood in primary forests, which provides nesting sites (Galbraith et al., 2019; Vázquez et al., 2011). Forests can shelter pollinators from climatic extremes, such as high temperatures, drought and strong winds, thereby buffering some of the known impacts of climate change (Dover et al., 1997; Ganuza et al., 2022; McDermott Long et al., 2016). Therefore, pollinators in general, including species that forage in open habitats as adults, may require access to multiple habitat types including but not limited to forests, to persist. Many studies have found a positive link between pollinator diversity, visitations, and forest cover or proximity to forests within predominantly agroecosystem landscapes (González Chaves et al., 2020; Marini et al., 2012a; Proesmans et al., 2019). Thus, maintaining forested areas around agroecosystems may enhance wild pollinator populations and potentially maximise the ecosystem services they provide (Kennedy et al., 2013). Despite abundant evidence that land-use change reduces pollinator diversity and services, crop- and region-specific studies that explicitly link forest cover to pollinator richness and visits remain scarce.

In the wake of global warming, increasing temperatures pose significant risks to agricultural crops. Tropical forests can have a significant cooling effect thereby potentially buffering agricultural crops from rising temperatures (Li et al., 2015). With ongoing climate change, open habitats such as agricultural lands experience higher maximum temperatures, lower minimum temperatures, and greater seasonal and interannual variability compared to forested habitats (de Frenne et al., 2019; Ewers & Banks-Leite, 2013; Von Arx et al., 2013), which may have significant consequences for pollinators (Wagner, 2020). Climate warming can affect pollinators in two ways: (a) directly, by imposing physiological constraints on pollinators thereby affecting their foraging activity with consequences for their reproduction and life span (Kalin et al., 1985; Scaven & Rafferty, 2013), and (b) indirectly, by altering resource availability. Increased temperatures can impact flower abundance (Anderson, 2016; Byers & Chang, 2017; Takkis et al., 2015), thereby indirectly affecting pollinator communities (Potts et al., 2006, 2010). Moreover, differential thermal preferences for visiting flowers may allow pollinators to minimize competition. Therefore, it is critical to understand foraging patterns of pollinators with respect to temperature to better understand the potential impacts of changing climate. Studies investigating the role of climate warming on plant-pollinator interactions have largely focused on phenological mismatches between flowering events and pollinator activity (see Hegland et al., 2009; Rafferty, 2017). There have been relatively few studies that investigate the impact of climate on the temporal activity of pollinators and its potential effects on pollination.

Much of the existing knowledge on pollination biology comes from temperate crop systems (Bentrup et al., 2019, 2021; Staton et al., 2019), whereas studies on tropical crops, especially in the Paleotropics, remain limited despite their high dependence on animal pollinators (Vizentin-Bugoni et al., 2018). Further, research in the context of agroforestry systems has focussed on coffee (Centeno-Alvarado et al., 2024; Jha & Vandermeer, 2010; Klein et al., 2002), where often native shade trees accompany the crop. Far fewer studies have focused on crops such as cashew, which are structurally simple and more open, and therefore may experience stronger impacts of land-use change and climate.

The associations between forest cover and climate on pollinator diversity are particularly relevant for cashew (*Anacardium occidentale*), because it relies on animal pollination for reproduction and fruit set (Freitas & Paxton, 1996; Vanitha & Raviprasad, 2019). Cashew, native to Brazil, was introduced in India by the Portuguese in the 16^th^ century (Kalaivanan, 2012), and India is now among the leading cashew nut exporters (Rege and Lee, 2022). The cashew-growing region of Maharashtra (MH) state, which is part of the Western Ghats-Sri Lanka Biodiversity Hotspot, has seen a 15% increase in cashew cultivation over the past decade (Chhaya et al., 2025), increasingly in monoculture form (Rege et al., 2022; Rege & Lee, 2022), underscoring the severity of land-use change in this region. Though the effects of remnant forests and climate change on pollinators in human-dominated landscapes are well-documented globally, few studies have disentangled their independent contributions simultaneously (but see Ganuza et al., 2022), an issue we address in this study. In particular, whether forest proximity mediates temperature in orchards, and the association of pollinator activity with temperature is an aspect that remains relatively understudied in the tree crop plantations context. Investigating how pollinators respond to habitat and temperature changes in the northern Western Ghats is critical, since the livelihoods of a significant section of the local population depend on the income from cashew farming (Nayak & Paled, 2018).

The aim of our study was to investigate the effect of landscape- and site-level factors on pollinator diversity and visits in a cashew-dominated landscape within India’s Western Ghats biodiversity hotspot. Specifically, we aimed to: (1) quantify the effect of landscape-level forest proximity (proportion of forest cover within 500 m buffer and distance to nearest forest patch) on local ambient temperature in cashew orchards, (2) assess the effects of forest proximity and temperature on pollinator richness, number of pollinators, number of visits, and species occurrences, while accounting for flower abundance.

## 2. Methods

### 2.1. Study area

The study was conducted during the dry season (January–April 2025) in cashew orchards in Dodamarg Taluk, Sindhudurg District, Maharashtra, India (Fig. S1 in Supplementary Material), coinciding with peak cashew flowering in the region (Fig. 1A; Madhu et al., 2025b). This landscape lies in the northern part of the Western Ghats–Sri Lanka Biodiversity Hotspot. The region receives an average annual rainfall of ∼3,500 mm, and the annual temperatures range between 12–46.5 (this work; Munje & Kumar, 2022). Cashew is the most important cash crop in the low-elevation areas of the region (Rege et al., 2022; Rege & Lee, 2022). The area under cashew cultivation is expanding rapidly (Chhaya et al., 2025; Rege et al., 2022), and nearly one-third of Sawantwadi and Dodamarg *Talukas* (sub-divisions of Sindhudurg district) are now under cashew (Rege & Lee, 2022). Previous studies in the region have found that cashew orchards negatively impact birds (Biswas et al., 2025; Madhu et al., 2025a), reptiles (Jithin et al., 2023), and amphibians (Jithin et al., 2025; Lad et al., 2025). In contrast, the presence of native trees within cashew orchards and surrounding forest cover has been shown to positively influence bird diversity (Madhu et al., 2025a).

**Figure 1.**
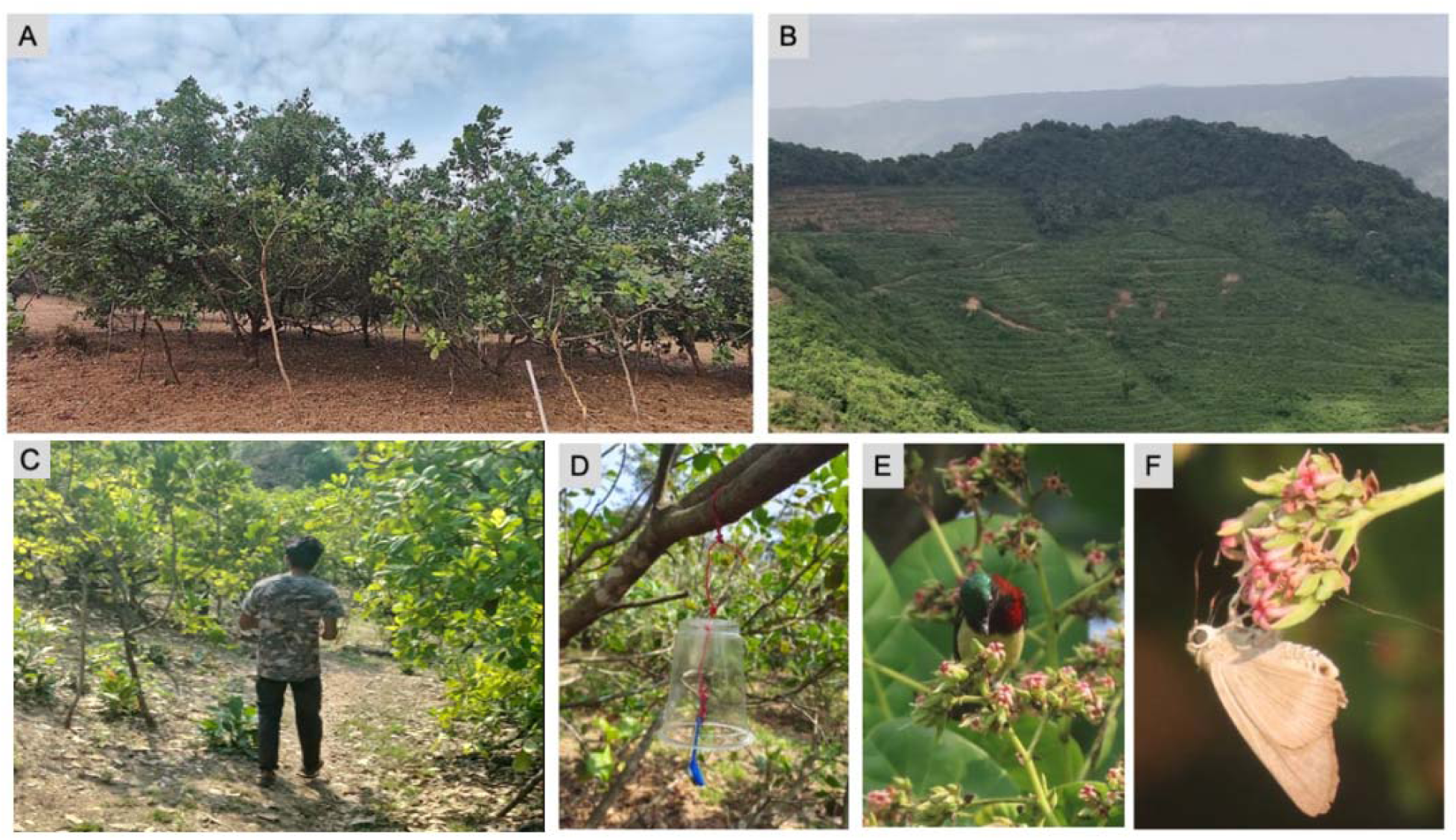
A) Typical cashew monoculture orchard at the study site, B) Cashew orchard adjoining a forest patch, C) AS sampling pollinators in a cashew orchard; image obtained with consent D) iButton deployed in cashew orchard for recording temperature, E) Crimson-backed Sunbird (*Leptocoma minima*), endemic species to the Western Ghats feeding on cashew nectar, and F) Brown Awl (*Badamia exclamationis*) feeding on cashew nectar. Photographs: Rohit Naniwadekar (A, B), Vishal Sadekar (C), Nandita Madhu (D) and Aditya Satish (E, F).

Forests are interspersed among cashew monocultures (Fig. 1B) and are either privately-owned or government-owned Reserved Forests. Privately-owned forests have been rapidly converted to cashew orchards (Fig. 1), and there are no protected areas at low-elevations in the region. The dominant natural vegetation type in low elevations is moist deciduous forest, with patches of evergreen forest interspersed. These forests harbour a mix of evergreen (e.g., *Syzygium* spp., *Knema attenuata*, *Machilus macrantha*, *Beilschmiedia dalzelli*) and deciduous tree species (e.g., *Terminalia paniculata*, *Terminalia elliptica*, *Careya arborea*).

### 2.2. Field methods

#### 2.2.1. Pollinator sampling

We (Aditya Satish and Vishal Sadekar) sampled for pollinators in 30 cashew orchards in the low-elevation areas of Dodamarg Taluk (mean elevation: 101.2 m; range: 34–281 m a.s.l.; Fig. 1; Fig. S1). We chose cashew orchards to represent variation in surrounding forest cover and distance to the nearest forest patch, based on previous studies (Biswas et al., 2025; Madhu et al., 2025a) and local ecological knowledge. We sampled for pollinators between 0730 to 1230 hr, coinciding with peak pollinator activity. Each survey consisted of seven 30-minute observation sessions with a 15-minute break between each session. Thus, we surveyed each orchard once, consisting of seven sessions per day, which totalled to 210 sessions for 30 orchards (3.5 h per orchard). During each session, two observers walked slowly around the orchard, recording individual animals that visited cashew flowers (Fig. 1C). Since the individuals interacted with the reproductive parts of cashew flowers, we refer to them as pollinators in this study. For each pollinator, we noted its species identity (or morphospecies when species-level identification was not possible), the number of visits made to flowers, and the nature of the interaction. The major pollinators (whose visits constituted 80% of the total visits) comprised a small set of birds, butterflies, and honeybees with well-resolved taxonomy, allowing reliable field identification (see Table S1 in Supplementary Material). Pollinators that could not be identified to species-level were classified to genus, tribe, or family, but were distinguished from similar-looking taxa by observable morphological traits and were recorded relatively infrequently. A visit was defined as a single contact of the pollinator with a flower’s reproductive parts, regardless of whether multiple contacts occurred on the same flower. Once a pollinator was encountered, one observer followed the individual until it was out of sight to record its visit frequency, while the other continued scanning for new pollinators. If a focal individual moved out of sight, and was later re-encountered during the observation session, it was treated as a separate individual. Since we consistently followed this protocol in all our orchards, we believe that any resulting biases were minimised. With this data (210 sessions across 30 orchards), we quantified pollinator richness, number of pollinators, and number of visits made by individual pollinators to cashew flowers.

#### 2.2.2. Quantification of flower abundance

Following pollinator sampling, we estimated floral abundance in each orchard by counting the number of panicles on ≥ 10 cashew trees. A reference tree was selected haphazardly, and the nearest trees (∼6 m apart) were subsequently chosen in a circular arrangement around it in each orchard. The canopy of each tree was divided into four quadrants, and panicles were visually counted in each quadrant and summed to obtain the tree-level panicle abundance. The mean panicle abundance across sampled trees was used as an index of flower abundance for each orchard.

#### 2.2.3. Temperature sampling

We deployed iButton temperature sensors (model: DS1921G-F5) to record ambient temperature in the orchards during pollinator sampling. In each orchard, ten iButton units were suspended ∼1.5 m off the ground from branches of the same ten trees used for flower abundance estimation (Fig. 1D). The iButtons were deployed for 24 hours, beginning the day before pollinator sampling and ending immediately after sampling was completed. Each iButton recorded temperature at 10-minute intervals. At the end of each sampling day, data were downloaded and the units were reset for deployment in the next orchard. Prior to the study, we tested for measurement error by suspending all iButton sensors at the same location for 24 hours and comparing their readings. As no detectable deviation was observed among sensors, we are confident that temperature measurement error was minimal.

#### 2.2.4. Calculation of forest cover and distance to forest

Using an existing land-use land-cover raster from a recent study (Rege et al., 2022), we calculated the proportion of forest cover around each sampling point within a predefined buffer radius, using the ‘landscapemetrics’, ‘raster’, and ‘sf’ packages in R (Hesselbarth et al., 2025; Hijmans et al., 2025; Pebesma et al., 2025). We selected a buffer radius of 500 m around each orchard, guided by its ecological relevance to pollinator foraging behavior and by consistency with previous landscape ecology studies (Campbell et al., 2022; Ganuza et al., 2022; Luppi et al., 2018; Marini et al., 2012b). In our study area, the primary pollinators of cashew flowers are small nectarivorous sunbirds (Family: Nectariniidae) (Fig. 1E), butterflies (Order: Lepidoptera) (Fig. 1F), bees and wasps (Order: Hymenoptera), and flies (Order: Diptera) (this work). Studies on related species have shown that they typically occupy small territories and forage over short distances, generally not exceeding 500 m (Araújo et al., 2004; Cant et al., 2005; Punchihewa et al., 1985; Zurbuchen et al., 2010). Therefore, the designated 500 m buffer size is likely to effectively capture the landscape most relevant to the foraging activities of cashew pollinators.

Using the same classified raster, we extracted the distance from each orchard to the nearest forest patch in QGIS (version 3.26.2).

### 2.3. Analysis

#### 2.3.1. Ambient temperature

We fit a linear mixed-effects model (Gaussian error) with forest cover (500 m), distance to nearest forest patch, and Julian day as fixed effects, and a random intercept for orchard identity, to assess the potential of macroscale temperature-regulating role of forests on cashew orchards. Since temperature progressively increases from January, we included Julian day as a predictor variable. The temperature values for each orchard were averaged for each of the 10 iButtons. We only considered the temperature data that coincided with the pollinator sampling duration (730 to 1230 hr) for this analysis. We assigned orchard ID as the random effect for the model.

#### 2.3.2. Pollinator richness, number of pollinators and number of visits

We used a generalized linear mixed model (GLMM) with a negative binomial distribution to model pollinator richness, number of pollinators, and number of visits as functions of distance to forest, forest cover, flower abundance, and temperature (both linear and quadratic terms). We included a quadratic term to account for potential non-monotonic relationships between response variables and temperature. Orchard ID was included as a random intercept to account for repeated measures across different sessions within the same orchard.

#### 2.3.3. Species responses

We used the Hierarchical Modeling of Species Communities (HMSC) framework (Ovaskainen & Abrego, 2020) to determine species responses to environmental predictors. This analysis complemented diversity and composition analyses and allowed us to determine species-specific responses to different environmental variables. We excluded rare species (59 out of the 66 species that occurred in less than 5% of sampling points; Table S1), as they provide little information on community assembly processes and can hinder Markov Chain Monte Carlo (MCMC) convergence (Ovaskainen & Abrego, 2020). We used the R package ‘Hmsc’ to fit the model in a Bayesian framework using default prior distributions (Tikhonov et al., 2025). We modelled species occurrence (with a probit link) as a function of (1) proportion of forest cover in the landscape (500 m buffer), (2) distance to the nearest forest patch, (3) flower abundance, and (4) temperature (linear and quadratic terms). We included the location of each sampling point and session ID as random effects to account for spatial and temporal autocorrelation. We sampled posterior distributions with three MCMC chains, thinned by 2000, yielding 250 samples for each chain. The first 100,000 samples were removed as burn-in. For all of the above analyses, the predictors were scaled to allow direct comparison across coefficients thereby determining their relative effects on response variables.

## 3. Results

### 3.1. Ambient temperature

The mean (± SD, range) temperature across the orchards during the sampling duration was 25.8 (± 1.4; 14–46.5). We did not find a statistically significant relationship between ambient temperature in orchards and forest cover, distance to the nearest forest patch, and Julian day (Table S2).

### 3.2. Pollinator richness, number of pollinators and number of visits

We observed 66 species visiting cashew flowers during the study, including 58 insect species and eight bird species (Table S1). In total, we observed 693 individuals (371 insects and 322 birds) interacting with cashew flowers (mean ± SD = 3.3 ± 4.0 individuals per session). These individuals made 9,714 visits in total (5,428 insect visits and 4,286 bird visits) (mean ± SD = 46.3 ± 132.0 visits per session) (Fig. S2).

### 3.3. Species richness

A generalized linear mixed model (GLMM) with a negative-binomial distribution revealed that flower abundance, temperature, and forest cover were the strongest predictors of species richness across sites (Table S3; Fig. 2). The first order quadratic term for temperature had a significant positive effect (coefficient = 0.401, *p* < 0.001), while the second-order quadratic term was not statistically significant (coefficient = −0.127, *p* = 0.051), suggesting that species richness increased monotonically with temperature (Fig. 2c; Table S3). Species richness also increased with forest cover within a 500 m radius around orchards (coefficient = 0.184, *p* = 0.05; Fig. 2a). As expected, flower abundance had a significant positive effect on pollinator richness (coefficient = 0.602, *p* < 0.001; Fig. 2b). Distance to the nearest forest patch had no significant effect (*p* = 0.605). The model explained a substantial portion of the variation in pollinator species richness, with a marginal *R²* (variance explained by fixed effects) of 0.48.

**Figure 2.**
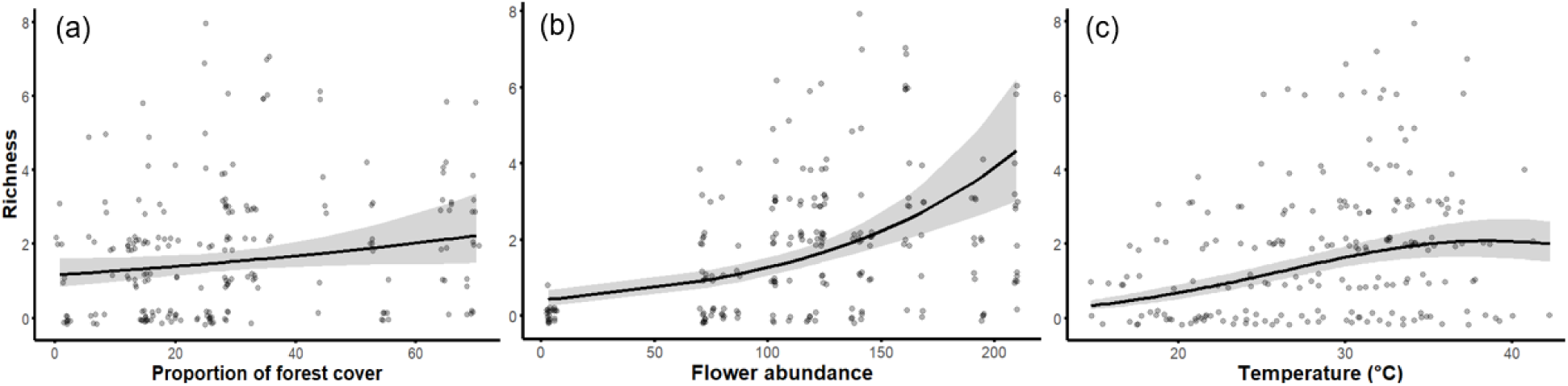
Predicted effects of select predictor variables on total pollinator richness per session in cashew orchards, based on negative-binomial GLMMs. Predicted relationships are shown for richness against (a) proportion of forest cover, (b) flower abundance (mean number of panicles on a cashew tree per orchard), and (c) temperature (°C, including both linear and quadratic terms). Observed data are shown as jittered grey points.

### 3.4. Number of pollinators

The number of pollinators was significantly influenced by flower abundance, forest cover, and temperature (Table S4; Fig. 3). The number of pollinators exhibited a hump-shaped relationship with temperature: the second-order quadratic term was significantly negative (coefficient = – 0.224, *p* = 0.006), indicating a peak around ∼30°C followed by a gradual decline at higher temperatures (Fig. 3c). Forest cover also had a significant positive effect (coefficient = 0.224, *p* = 0.032), with predicted pollinator numbers increasing across the forest cover gradient (Fig. 3a).

**Figure 3.**
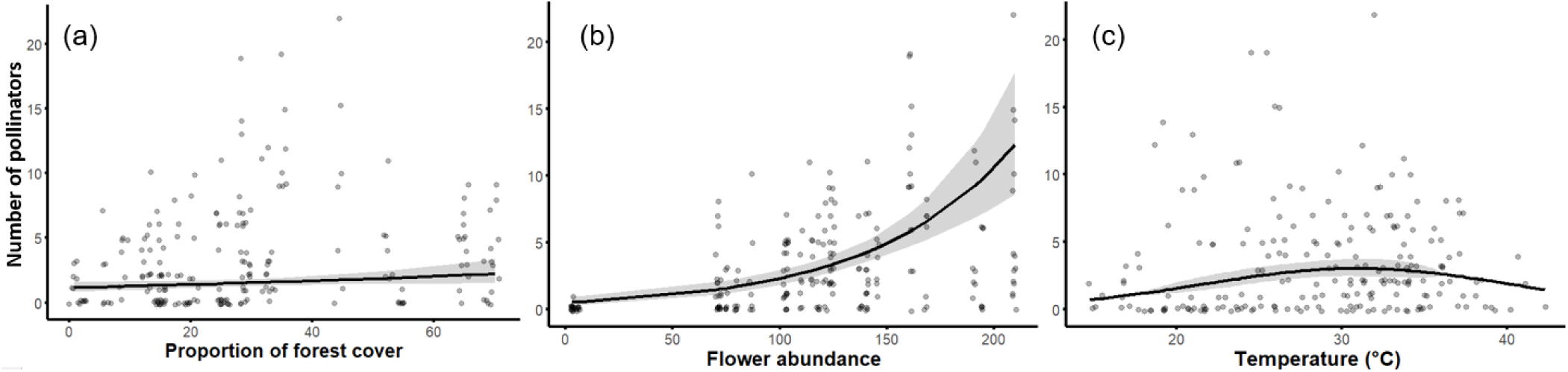
Predicted effects of select predictor variables on total number of pollinators per session in cashew orchards, based on negative-binomial GLMMs. Predicted relationships for number of pollinators are shown against (a) proportion of forest cover, (b) flower abundance (mean number of panicles per tree per orchard), and (c) temperature (°C, including both linear and quadratic terms). Observed data are shown as jittered grey points.

Flower abundance had a significant positive effect on the number of pollinators (coefficient = 0.823, *p* < 0.001), with a steep increase at higher flower densities (Fig. 3b). Distance to the nearest forest patch had no significant effect (*p* = 0.807). The model explained a considerable amount of variation in the data, with a marginal *R^2^*of 0.507.

### 3.5. Number of visits

The number of visits was strongly influenced by temperature, forest cover, and flower abundance (Table S5; Fig. 4). The second-order quadratic term for temperature was significantly negative (coefficient = –0.41, *p* = 0.001), suggesting the hump-shaped response observed in the predicted relationship (Fig. 4c), with visits peaking around 30°C and declining thereafter. Forest cover also showed a significant positive effect (coefficient = 0.531, *p* = 0.002), and the predicted plot indicated an increase in visits with increasing forest cover (Fig. 4a). Flower abundance had a significant positive effect (coefficient = 1.388, *p* < 0.001), with a sharp exponential increase in the number of flowers at higher flower densities (Fig. 4b). Distance to the nearest forest patch had no significant effect (*p* = 0.41). The model explained substantial variation in the number of visits, with a marginal *R^2^* of 0.598 for fixed effects. Bird visits to cashew flowers peaked during the second sampling session (0815–0845 hr; mean temperature: 22.3□), while insect visits peaked during the fifth sampling session (1030–1100 hr; mean temperature: 32.6□) (Fig. 4d).

**Figure 4.**
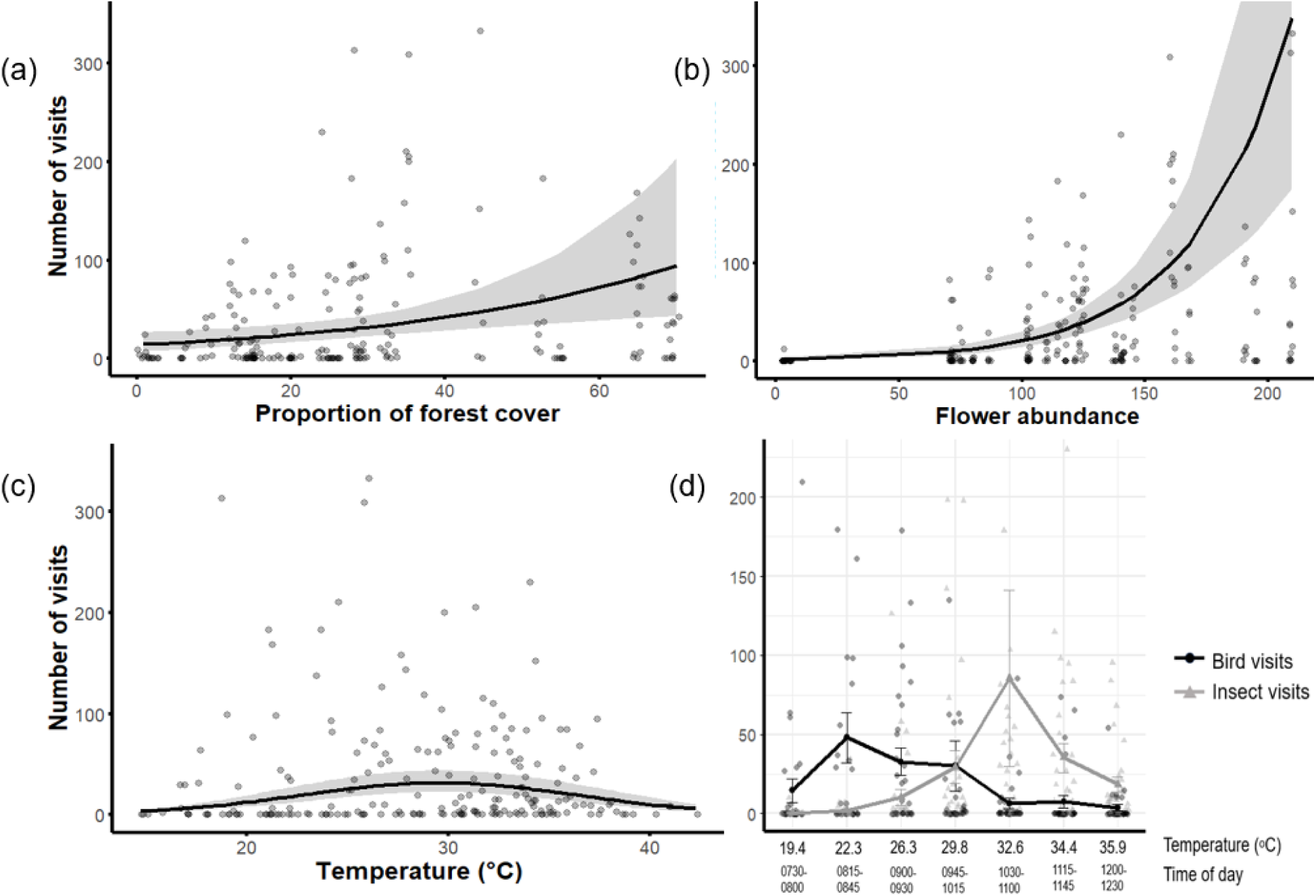
Predicted relationships between select predictor variables and the total number of visit by pollinators per session in cashew orchards, based on negative-binomial GLMMs. Predicted visits are shown as a function of (a) proportion of forest cover, (b) flower abundance (mean number of panicles per tree per orchard), and (c) temperature (°C, including first and second order terms). Y-axes are limited to 350 visits for clarity; two extreme values (475 and 1700 visits) were included in the models but not displayed here. Panel d shows variation in the mean number of visits to cashew flowers by birds and insects across seven time-of-day sessions and associated temperatures. Lines connect session-wise means (± 1 SE), with error bars indicating standard errors. The average temperature recorded during each session is shown on the x-axis above the corresponding time block. A few extreme outliers (n = 4; values >250) were excluded from the plot for visual clarity but were retained in the calculation of means and standard errors. Observed data are shown as jittered grey points in all panels.

### 3.6. Species responses

Species-level responses to environmental variables varied in both direction and strength (Fig. 5a). Temperature (42.7%) explained the bulk of the variation in the probability of occurrence of pollinators, followed by flower abundance (22.3%). Spatial (8.1%) and temporal (9.3%) random effects explained relatively little variation in the data. For most pollinators, except *Apis cerana* and *Delias eucharis*, occurrence probability showed a hump-shaped relationship with temperature. *Apis cerana* occurrence showed an increasing trend with temperature. Flower abundance had a significant positive effect on occurrence probability for all species. Forest cover positively influenced *Badamia exclamationis*, while the occurrence of *Euploea core* and *Delias eucharis* declined with increasing distance to the nearest forest patch. These results highlight species-specific ecological preferences and differential sensitivity to habitat and climatic gradients. Variance partitioning revealed that the relative contribution of predictor variables varied across species (Fig. 5b). Tjur *R^2^* values indicated variable model performance across species, with *Leptocoma minima* (Tjur *R^2^*= 0.47) and *Cinnyris asiaticus* (Tjur *R^2^* = 0.41) showing the highest values, indicating that their occurrences were better explained by the model.

**Figure 5.**
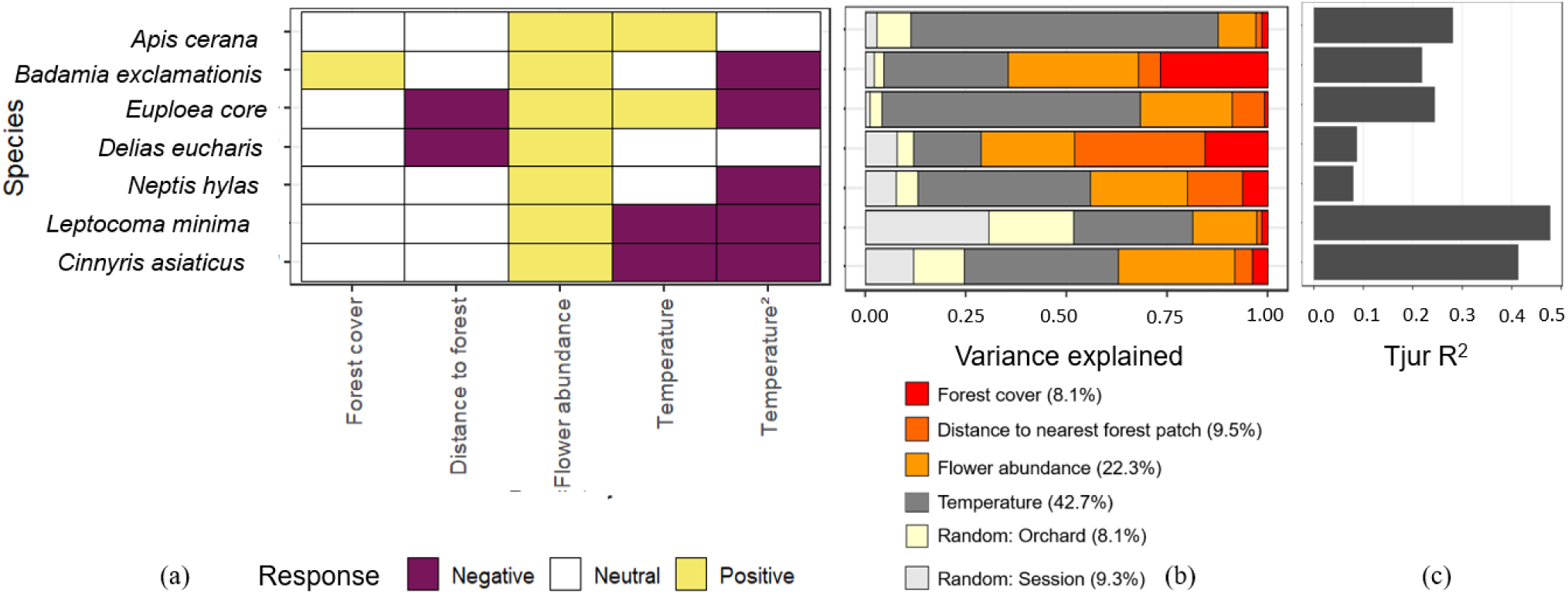
Species-level responses for seven species: *Apis cerana*, *Badamia exclamationis*, *Euploea core*, *Delias eucharis*, *Neptis hylas*, *Leptocoma minima*, and *Cinnyris asiaticus*. Composite figure showing (a) the beta plot (posterior regression parameters), (b) variance partitioning plot, and (c) the Tjur *R^2^* plot. The beta plot displays species responses to predictors. Yellow and violet boxes indicate statistically significant (posterior support ≥ 95%) positive and negative responses, respectively. The variance partitioning plot shows the proportion of variance in species occurrence explained by different predictors. The Tjur *R^2^*plot shows the overall variation in species occurrences explained by the model.

*Apis cerana* (Tjur *R^2^* = 0.28), *Badamia exclamationis* (Tjur *R^2^* = 0.21), and *Euploea core* (Tjur *R^2^* = 0.24) had moderate values, while the model performed least well for *Delias eucharis* and *Neptis hylas* (Tjur *R^2^* = 0.08 for both), where explanatory power was relatively low (Fig. 5c).

## 4. Discussion

We tested the effects of site-(flower abundance, temperature) and landscape-level (proportion of forest cover, distance to nearest forest patch) factors on cashew pollinator richness, number of pollinators, number of visits by each pollinator, community composition and individual species occurrences. At the community level, both site- and landscape-level factors positively influenced pollinator richness, number of pollinators, and number of visits made by pollinators.

Additionally, the number of pollinators and their visits peaked within a relatively narrow temperature range (∼28-32°C). Insects and birds differed in their visitation patterns on flowers, corresponding with temperature changes, likely reflecting underlying physiological differences. At the species level, temperature and flower abundance explained most of the variation in species occurrences. Forest cover positively influenced *Badamia exclamationis*, while other lepidopterans such as *Delias eucharis* and *Euploea core* were more frequently recorded in cashew orchards near forest patches. Occurrence of all species were positively associated with flower abundance. Temperature effects were species-specific and appear to reflect ecological differences in thermal tolerance and foraging activity.

### 4.1. Role of forest patches

#### 4.1.1. Effects of forest proximity on pollinators

After accounting for the effects of flower abundance in our models, we find significant effects of both landscape-level factors like forest cover and distance and temperature. While forest cover positively influenced community-level metrics such as the pollinator richness, number of pollinators, and number of visits of each pollinator – species-level responses of lepidopterans were influenced by both forest cover and distance to the forest. This finding is well-supported by multiple studies in the literature (Ulyshen et al., 2023). The greater number of visits in orchards in proximity to forests could be because of spillover effects from nearby forests that may contain a greater amount of resources (Urban-Mead et al., 2022). Moreover, forests benefit pollinators by providing refuge, nesting, and food resources (Proesmans et al., 2019; Ulyshen et al., 2023).

Lepidopterans may benefit from surrounding forests because forests often contain their larval host plants, thereby maintaining population sizes (Dennis-Macfie et al., 2003). We observed higher occurrences of *Badamia exclamationis* in cashew orchards with ambient forest cover, and higher occurrences of *Delias eucharis* and *Euploea core* in orchards closer to forest (Figure 6a). The larval hostplants of *Badamia exclamationis* (*Terminalia* spp.), *Delias eucharis* (*Ficus* spp., *Tabernaemontana alternifolia*, *Holarrhena pubescens* and *Wrightia tinctoria*) and *Euploea core* (*Dendrophthoe* spp. and *Butea monosperma*) (Nitin et al., 2018) are common in the forests of the northern Western Ghats surrounding the cashew orchards, which may help explain their associations with forest. The higher number of visits in orchards with greater surrounding forest cover may directly translate to greater yields that could be beneficial for local farmers (Freitas et al., 2014; Kapoor & Sharma, 2022), but this needs further investigation in our study area.

Interestingly, the occurrences of the top three pollinators (*Leptocoma minima*, *Cinnyris asiaticus*, and *Apis cerana*) were not influenced by either forest cover or distance to forest.

*Cinnyris asiaticus* prefers open habitats and its occurrence was negatively associated with forest cover (Biswas et al., 2025; Madhu et al., 2025a). Evidence regarding the open-habitat preference for *Leptocoma minima* is mixed: Biswas et al., 2025 found that its occurrence was lower in cashew orchards, but Madhu et al., 2025a reported a positive association between its probability of occurrence and cashew cover at large scales in the same study area. A similar mixed trend is observed for *Apis cerana* (Klein et al., 2002; Koetz, 2013), with no clear consensus on whether it is more associated with forests or open habitats. These could be because *A. cerana* nest in cavities that are perhaps common in forested habitats (Corlett, 2004; Koetz, 2013; Seeley et al., 1982), but forages in open agricultural systems which seasonally offer abundant food resources. We do not know of any farmers in our study area that maintain domesticated *Apis cerana* colonies (Vishal Sadekar, pers. obs.), and therefore we assume that all our observations of this species are of wild individuals. Since the occurrence of the three common pollinators was not associated with forest cover and distance to forests, the observed community-level patterns are a likely outcome of cumulative effects of all the other species in the community.

Although cashew orchards may also offer pollinators a reliable and abundant food source, these resources are available only for a short period of time (Fig. S3a). Madhu et al., 2025b found that while the flower abundance was higher in cashew orchards and agroforests during peak cashew flowering season, the decline in bird-visited floral resources was also significantly faster over time compared to adjoining forest habitats. Forest cover around orchards may provide pollinators with additional floral and non-floral resources during and beyond the crop flowering season, which can increase their propensity to occur around orchards with sufficient surrounding forest cover (Dover & Settele, 2009; Kremen et al., 2002; Öckinger & Smith, 2007). We also observed that soils in the cashew orchards of our study area tend to be less acidic (median pH = 7.2 in cashew and 6.5 in forests) and poor in potassium, phosphorus and other minerals (Ca, Fe, Mn, Zn, among others) compared to forests (Sadekar et al., 2025; Fig. S4), indicative of intensive pesticide use that may be harmful to pollinators (Basu et al., 2024; Uhl & Brühl, 2019). Natural forests around intensively cultivated croplands can buffer the negative impacts of such chemical inputs on wild pollinator communities (Park et al., 2015).

### 4.2. Effects of temperature on pollinators

#### 4.2.1 Direct effects

We demonstrate the important role that temperature plays in mediating the richness, numbers and occurrence of pollinators, and number of cashew flowers visited. Birds, which are warm-blooded animals, visited the flowers significantly earlier compared to insect pollinators. The richness of pollinators peaked at around 35, while the number of pollinators that we detected on flowers peaked at around 29. The peak in overall richness at a higher temperature may be due to higher richness of insects (58 species) that tend to visit flowers at higher temperatures, which far outnumbered the richness of bird pollinators (8 species) that largely visited flowers at lower temperatures earlier during the day.

Among the relatively common pollinators, each has its own preferred temperature range suggesting that foraging activity for each pollinator species is constrained by a thermal optimum (Williams et al., 2007), above or below which it may not be energetically feasible to forage.

Under high ambient temperatures, only species able to tolerate thermal stress remain active, and this shapes when different pollinators can visit flowers (Goodwin et al., 2021; Vicens & Bosch, 2000; Willmer, 1983). The thermal limits of pollinators may dictate the timing and patterns of their foraging activity. The ongoing climate warming can lead to shifts in the time of day at which pollinators visit flowers (Willmer, 1983; Willmer & Stone, 2004) or restrict foraging trips to shorter distances or durations to avoid overheating during flight at elevated temperatures (Scaven & Rafferty, 2013). As a result, with climate change and rising temperatures, the window for optimal foraging is likely to become narrower, ultimately leading to fewer visits, shorter interaction times and earlier peaking of foraging activity (de Manincor et al., 2023; Williams et al., 2007). Such shifts in activity patterns may cause changes in the temporal composition of the pollinator community, which could have cascading effects on the timing and duration of flower visits, the movement of pollen across individual plants and the effectiveness of cashew pollination. Additionally, with higher temperatures pollinators may suffer population declines (Pyke et al., 2016) due to heat stress (Jevanandam et al., 2013), or shift to higher elevations in response to warming (Marshall et al., 2020). *Apis cerana* occurrence was positively associated with temperature, as this species was active during warmer parts of the day, a finding consistent with other studies examining pollination by *Apis* spp. (Celebrezze & Paton, 2004). In contrast, *Leptocoma minima* and *Cinnyris asiaticus* (family Nectariniidae), and lepidopterans *Badamia exclamationis* (family Hesperiidae), *Neptis hylas* (family Nymphalidae) and *Euploea core* (Nymphalidae) show unimodal patterns with temperature suggesting niche partitioning among pollinators.

#### 4.2.2. Indirect effects through flower abundance

Cashew requires warm, dry conditions to flower (Mir et al., 2014) and high temperatures beyond 34 are known to result in the desiccation of flowers (Fig S3c; Mir et al., 2014). We also observed a strong association between flower abundance and pollinators, a finding well-established in the literature and closely linked to nectar and pollen availability (Hegland & Boeke, 2006; Potts et al., 2003; Potts et al., 2006). This strong association between cashew flower abundance and pollinators makes them vulnerable to climate-mediated effects. Higher temperatures can reduce flower size and flowers per plant, thus reducing floral display size (Brunet & Van Etten, 2019; Descamps et al., 2021). Smaller floral displays may limit food availability for pollinators (Makino et al., 2007; Ohashi & Yahara, 2001), with feedback on pollinator population sizes (Häussler et al., 2017) and cashew pollination. Additionally with changing temperatures, there might be phenological mismatches between flowers and pollinators, further complicating the dynamics between plant-pollinator interactions (Gérard et al., 2020). Altogether, these factors make the impacts of climate on cashew pollination a multifaceted problem that warrants further investigation.

### 4.3 Effects of forest cover on temperature

Forests are well-known to provide cooling benefits relative to proximal open landscapes (de Frenne et al., 2019; Ewers & Banks-Leite, 2013). However, we failed to find support for this regulatory effect on cashew orchards in proximity to forests. Cashew orchards are open habitats with low tree density, short trees and sparse canopy cover (Madhu et al., 2025a). These conditions may negate the potential influence of macroclimatic buffering of forests. Moreover, in fragmented landscapes with relatively smaller forest patches, as in our study, this influence may be weak, reducing their capacity to buffer temperatures in cashew orchards. Few studies have simultaneously examined the effects of forest proximity and temperature on pollinators. In our study, the absence of a strong correlation between landscape factors (i.e. proportion of forest cover within a 500 m buffer and distance to nearest forest patch) and temperature allowed us to investigate the relative effects of these variables on pollinators.

### 4.4. Uncertain future of cashew

Taken together with the documented effects of land-use and temperature on pollinators, our findings suggest that the future of cashew production in the region under a business-as-usual scenario is uncertain. Cashew flowering depends on prolonged dry periods and relatively cool winter temperatures (Mir et al., 2014), but these conditions are increasingly threatened by rising temperatures under climate change (Fig. S3c). Because flower abundance strongly influences pollinator richness, the number of pollinators, and the number of visits, as demonstrated in this study, reduction in flowering is likely to cascade into reduced pollinator services. Moreover, the strong temperature associations for the most common pollinators indicate that climate-driven fluctuations in temperature will further depress visitation rates. At the same time, global evidence shows that land-use change reduces pollinator diversity and abundance (Ganuza et al., 2022; Millard et al., 2021; Ulyshen et al., 2023), and cashew orchards in our landscape are rapidly expanding at the expense of forests (Rege, Bodhankar Warnekar, et al., 2022). These landscape-level changes are therefore likely to compound the negative effects of climate on both pollinators and cashew flowering. Given that pollinator richness and abundance are known to strongly influence crop yields (Garibaldi et al., 2013), the combined pressures of climate and land-use change highlight a precarious future for cashew yields in the region if current trends continue.

### 4.5 Temporal niche differentiation between bird and insect pollinators

Our results indicate temporal niche partitioning among cashew pollinator communities. Birds (largely sunbirds) visited flowers during the early morning. In contrast, insect activity peaked at 1030-1100 hours when temperatures were much warmer (∼32°C), when bird visits were negligible. However, insect activity (except for *Apis cerana*) dropped after 1100 hours. Therefore, there are differences in visitation patterns between birds, which are endotherms and therefore able to regulate their body temperatures, and insect pollinators, which are ectotherms. This temporal partitioning potentially allows them to coexist, a pattern also reported in other studies (Celebrezze & Paton, 2004; Vaughton, 1996). Further, it appears that nectar volume and sugar production in cashew is minimal in the early morning from 0700-0900 hours, gradually increasing to a peak at 1300 hours (Bhattacharya, 2004), suggesting that the rewards gained by birds and insects are likely different. The decline in bird visits when faced with the rise in insect visits may also be influenced by potential competitive costs— there is evidence that indicates that nectarivorous birds may refrain from nectar-feeding later in the day, when the foraging activity of honeybees (the most common insect pollinator in our study) is at its peak and leads to reduced nectar availability, suggesting a degree of competitive exclusion as a consequence of exploitative competition may be at work (Hansen et al., 2002).

Our study is one of the few that uses a multi-taxa approach across unrelated taxonomic groups in a community ecology pollination study (as in Ganuza et al., 2022). Past studies have largely focused on a single taxonomic group, largely bees, given their dominance in pollination systems (Klein et al., 2006). However, the contribution of many non-bee taxa (lepidopterans, birds, dipterans etc.) to pollination remain understudied. This is especially important to correct, because different groups have varied life-history strategies and morphologies that shape their interactions with flowering plants differently. In our study, we were able to establish sunbirds (*Leptocoma minima* and *Cinnyris asiaticus*, family Nectariidae) as novel and regular vertebrate pollinators of cashew (Table S1), which has hitherto been largely thought of as a bee-pollinated species (Freitas et al., 2002; Freitas & Paxton, 1996; Vanitha & Raviprasad, 2019). We also report novel interactions between cashew and members of other insect groups, such as lepidopterans, non-bee hymenopterans, dipterans, coleopterans, and hemipterans (Table S1). However, the multi-taxa approach also has certain limitations. In our case, sunbirds are much larger and more conspicuous than *Apis cerana* and other insect pollinators, which may have introduced detectability biases in our sampling. We sought to minimize this effect by conducting observations from morning, when birds are most active, through the afternoon, when insect activity peaks, thereby capturing sufficient visitation data from both groups. Moreover, relatively few visitations on cashew flowers per session allowed us to carefully scan flowers for pollinators.

### 4.6. Future directions and conservation implications

Future studies can incorporate measurement of yield to quantify the impact of pollination services. If a positive relationship can be found, then it makes a strong case for increasing food, habitat and nesting resources of pollinators around agricultural areas, thus enhancing pollinator pollinations/movement in agroecosystems. For example, cashew orchard owners in collaboration with conservation organisations can explore (a) increasing the amount of specific floral and non-floral forage i.e., planting larval host plants of butterflies (Majewska & Altizer, 2020) or planting a diversity of visually attractive (native) flowering plants for pollinators that can draw them to orchards (Ghazoul, 2006) (b) retaining dead wood that can function as nesting sites for cavity-nesting bees (Simanonok et al., 2022), like *Apis cerana*, one of the chief pollinators in our study, and (c) enhancing opportunities for habitat connectivity between orchards with strips of native vegetation, that can allow movement of pollinators within heterogeneous landscapes (Steffan-Dewenter et al., 2002).

Since cashew farmers use pesticides in their orchards (Aditya Satish and Vishal Sadekar, pers. obs.), which is perhaps reflected in their soil composition (Fig S5), we also think that future studies should also explore the impact of pesticides on insect pollinators. We also recommend investigating the role of forests and agroforests in providing a buffer against heat extremes induced by climate change, relative to cashew monocultures, that can provide more support for restoration in forests that have been repeatedly disturbed in the landscape (Biswas et al., 2024). Given the temperature sensitivity of pollinator visits, future studies should investigate habitat suitability for cashew and its pollinators under changing climate to determine potential phenological mismatch (sensu Magrach & Ghazoul, 2015). Understanding mismatches and rewiring of interactions between cashew and its pollinators under future climate scenarios will be key to anticipating disruptions to pollination services and implementing adaptive management strategies for sustainable cashew production.

## CRediT authorship information statement

**Conceptualisation:** RN, AS, NM, VS, RaNa, AR; **Methodology:** AS, RN, VS; **Investigation:** AS, VS, NM; **Data curation:** AS; **Formal analysis:** AS, RN; **Funding acquisition:** RN; **Writing original draft:** AS, RN; **Writing review and editing:** All authors.

## Data Availability

All data files associated with the manuscript will be uploaded on Zenodo upon acceptance of the manuscript.

## Acknowledgements

We thank the Maharashtra Forest Department for the permission to conduct the study. We are grateful to Pravin Desai and family for assistance in the field. We would also like to thank Kruti Chhaya, Jithin Vijayan, Vijay Karthick, Anand Osuri, KP Akilan, and Bindu K for their valuable inputs, and cashew farmers of Dodamarg for allowing us to work in their orchards.

## Appendix A. Supplementary Information

**Figure S1.**
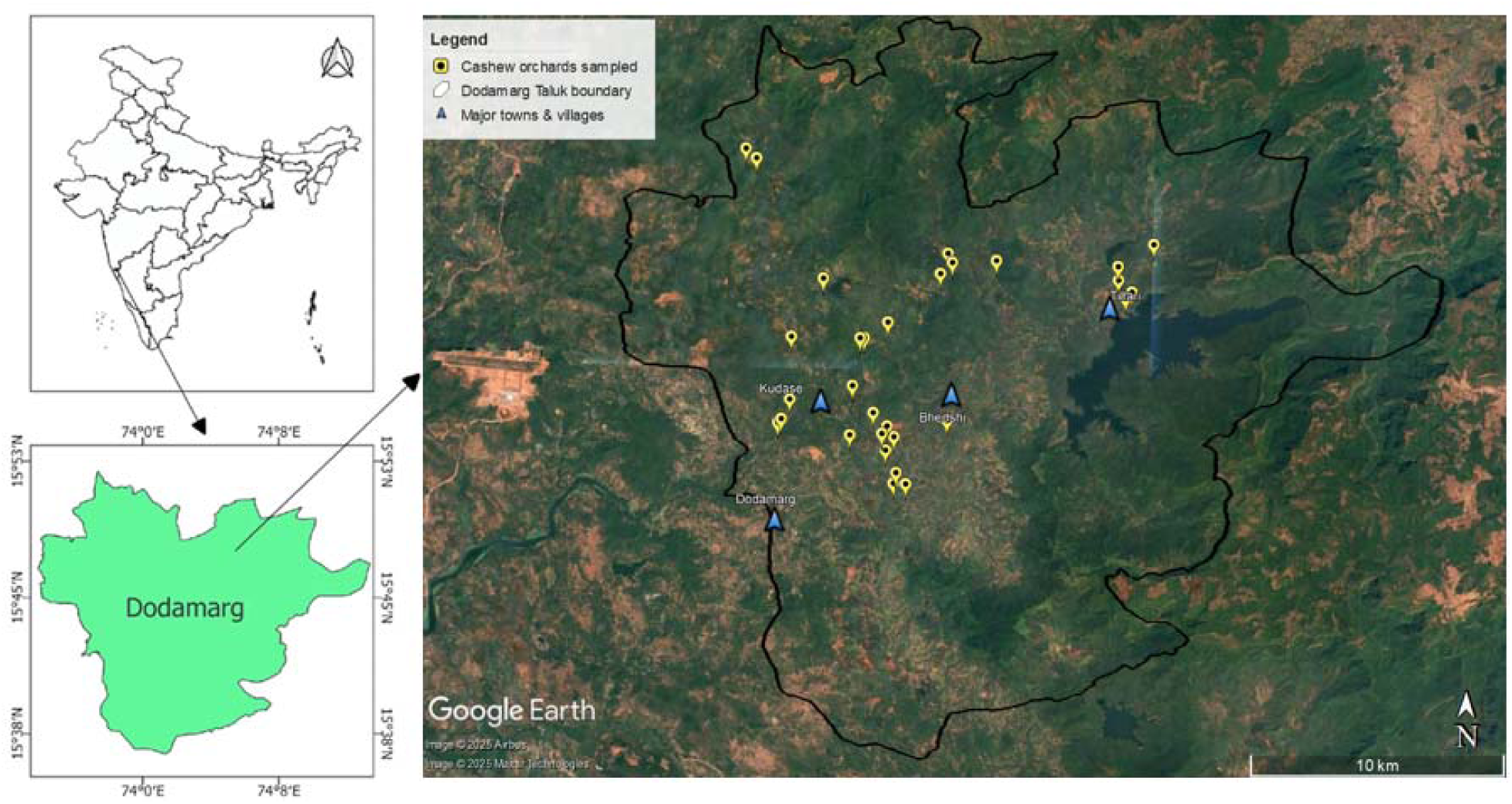
Study area. Map depicting the 30 sampled cashew orchards in Dodamarg Taluk, Maharashtra, India.

**Figure S2.**
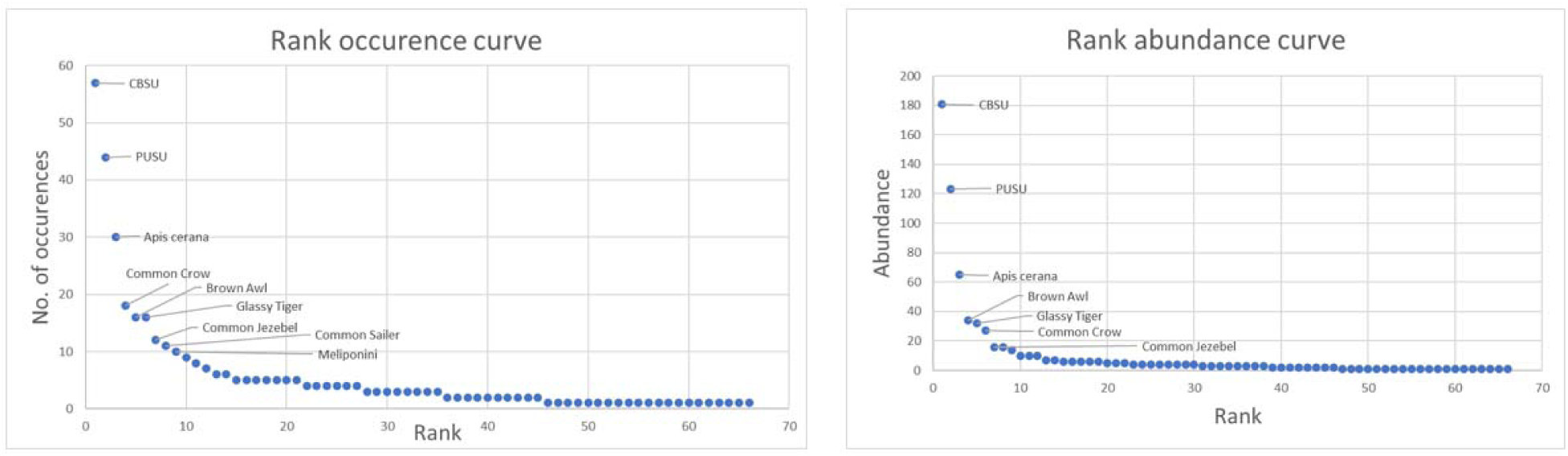
Rank distribution of taxa recorded visiting cashew flowers across all orchards. Rank occurrence curve showing the total number of occurrences of each species (left). Rank abundance curve showing the total number of individuals recorded per species across all sessions (right).

**Figure S3.**
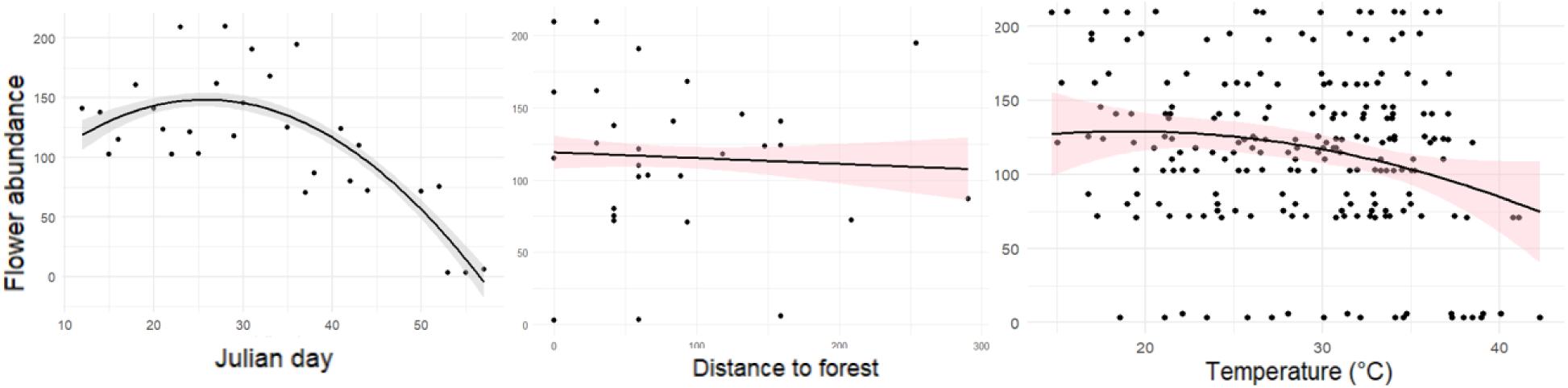
Relationships between flower abundance with Julian day (modelled with first order and second order quadratic terms) (left), distance to forest (centre) and temperature (modelled with first order and second order quadratic terms) (right).

**Figure S4.**
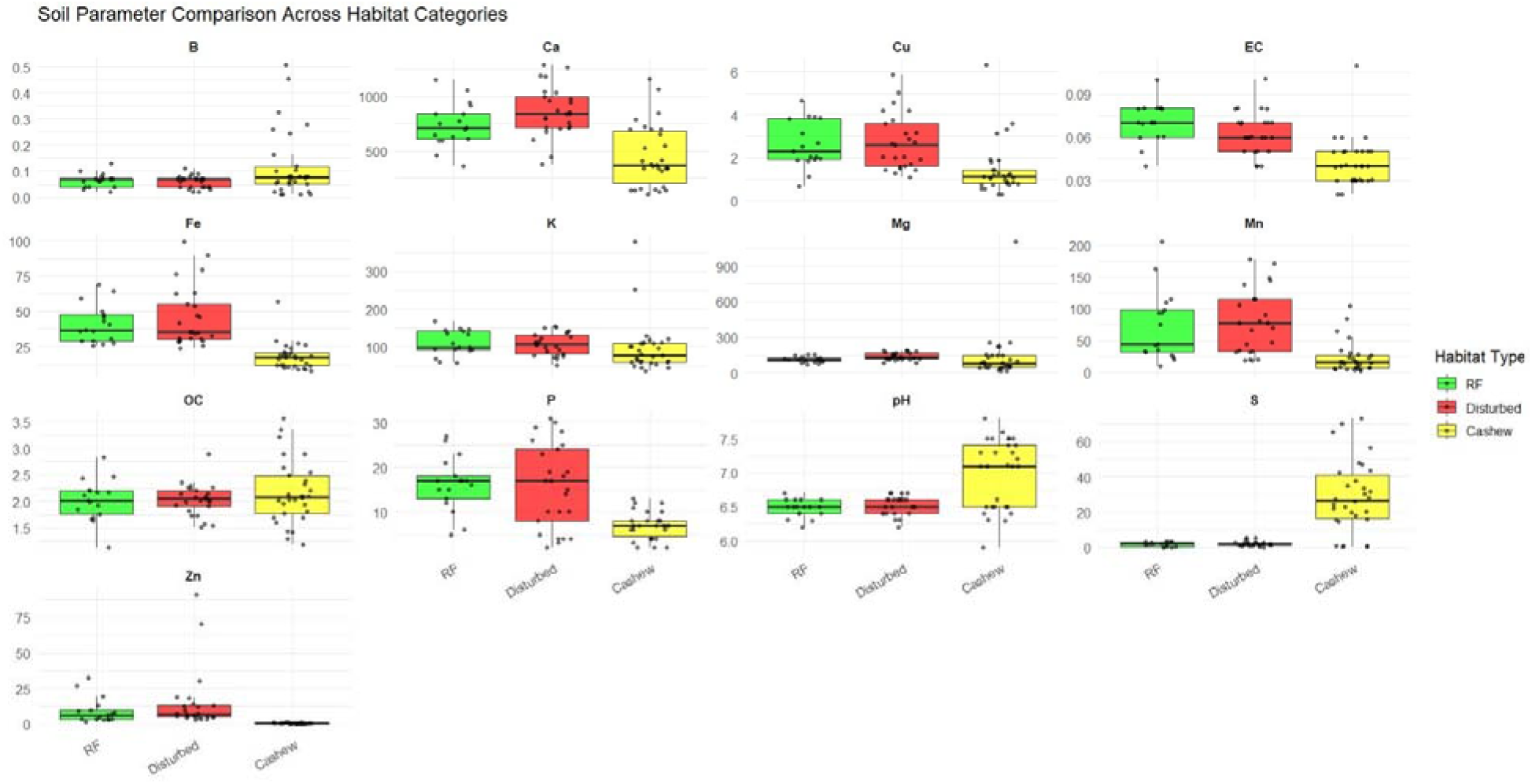
Comparison of soil parameters between RF (Reserve Forest), Disturbed forest and cashew orchards in Dodamarg, Maharashtra, India. The thirteen soil chemical properties we estimated are as follows: available boron (B), calcium (Ca), copper (Cu), electrical conductivity (EC) at 25°C, iron (Fe), potassium (K_2_O), magnesium (Mg), manganese (Mn), organic carbon (OC), available phosphorus (P_2_O_5_), pH, sulfur (S), and zinc (Zn).

**Table S1.** List of cashew pollinators. Some names correspond to morphospecies or those we could not identify in the field to the species-level.

| No. | Taxon | Species | Number of individuals recorded |
| --- | --- | --- | --- |
| 1 | Aves | *Leptocoma minima* | 181 |
| 2 | Aves | *Cinnyris asiaticus* | 123 |
| 3 | Insecta | *Apis cerana* | 68 |
| 4 | Insecta | *Badamia exclamationis* | 34 |
| 5 | Insecta | *Parantica aglea* | 32 |
| 6 | Insecta | *Euploea core* | 28 |
| 7 | Insecta | *Delias eucharis* | 17 |
| 8 | Insecta | *Neptis hylas* | 16 |
| 9 | Insecta | Meliponini | 14 |
| 10 | Insecta | Bombyliidae sp. 1 | 12 |
| 11 | Insecta | Wasp sp. 2 | 10 |
| 12 | Insecta | Bombyliidae sp. 2 | 10 |
| 13 | Insecta | Lycaenidae sp. | 7 |
| 14 | Insecta | *Appias albina* | 7 |
| 15 | Insecta | *Castalius rosimon* | 6 |
| 16 | Insecta | *Delta* sp. | 6 |
| 17 | Insecta | Diptera sp. | 6 |
| 18 | Insecta | *Apis florea* | 6 |
| 19 | Aves | *Dicaeum concolor* | 6 |
| 20 | Insecta | *Polistes* sp. | 6 |
| 21 | Insecta | Dolichopodidae sp. | 5 |
| 22 | Insecta | Syrphidae sp. | 5 |
| 23 | Insecta | Wasp sp. 1 | 5 |
| 24 | Insecta | *Neptis jumbah* | 4 |
| 25 | Insecta | Anthophila sp. | 4 |
| 26 | Insecta | *Xylocopa ruficornis* | 4 |
| 27 | Insecta | *Campsomeriella collaris* | 4 |
| 28 | Insecta | Scoliidae sp. | 4 |
| 29 | Aves | *Pycnonotus jocosus* | 4 |
| 30 | Insecta | *Junonia iphita* | 4 |
| 31 | Insecta | *Tirumala limniace* | 4 |
| 32 | Insecta | *Pantoporia* sp. | 3 |
| 33 | Insecta | *Dysphania* sp. | 3 |
| 34 | Insecta | *Tagiades gana* | 3 |
| 35 | Insecta | Nymphalidae sp. | 3 |
| 36 | Insecta | Bombyliidae sp. 3 | 3 |
| 37 | Insecta | Unidentified Insecta | 3 |
| 38 | Insecta | *Euthalia aconthea* | 3 |
| 39 | Insecta | *Catopsilia pomona* | 2 |
| 40 | Insecta | Eumeninae sp. | 2 |
| 41 | Insecta | Coleoptera sp. | 2 |
| 42 | Insecta | Halictidae sp. | 2 |
| 43 | Insecta | *Junonia lemonias* | 2 |
| 44 | Insecta | *Graphium teredon* | 2 |
| 45 | Insecta | Pieridae sp. | 2 |
| 46 | Insecta | *Xylocopa* sp. | 2 |
| 47 | Insecta | *Rapala manea* | 1 |
| 48 | Aves | *Pycnonotus cafer* | 1 |
| 49 | Insecta | Hemiptera sp. | 1 |
| 50 | Insecta | *Catopsilia pyranthe* | 1 |
| 51 | Insecta | *Tajuria cippus* | 1 |
| 52 | Insecta | *Ypthima huebneri* | 1 |
| 53 | Insecta | *Elymnias caudata* | 1 |
| 54 | Insecta | *Graphium* sp. | 1 |
| 55 | Insecta | *Ypthima baldus* | 1 |
| 56 | Insecta | *Graphium agamemnon* | 1 |
| 57 | Insecta | *Junonia atlites* | 1 |
| 58 | Insecta | *Junonia orithya* | 1 |
| 59 | Insecta | Red-bellied Scoliidae sp. | 1 |
| 60 | Insecta | *Catochrysops strabo* | 1 |
| 61 | Insecta | *Papilio polytes* | 1 |
| 62 | Aves | *Orthotomus sutorius* | 1 |
| 63 | Aves | *Prinia inornata* | 1 |
| 64 | Aves | *Aegithina tiphia* | 1 |
| 65 | Insecta | *Pachliopta aristolochiae* | 1 |
| 66 | Insecta | *Luthrodes pandava* | 1 |

**Table S2.** Output of the linear mixed-effects model (Gaussian error structure) that examined the relationship between ambient temperature and forest cover (500 m buffer), distance to nearest patch, and Julian days.

| Predictor | Estimate | SE | *t* | *p* |
| --- | --- | --- | --- | --- |
| Intercept | 25.755 | 0.264 | 97.395 | <0.001 |
| Forest cover | 0.087 | 0.275 | 0.316 | 0.754 |
| Distance patch | -0.133 | 0.277 | -0.479 | 0.636 |
| Julian days | 0.299 | 0.266 | 1.122 | 0.272 |
|  | **Marginal R<sup>2</sup>: 0.046** |  |  |  |
|  | **Conditional R<sup>2</sup>: 0.933** |  |  |  |
|  | **Random effect:** intercept for Orchard ID (n = 30 orchards), SD = 1.44 |  |  |  |

**Table S3.**
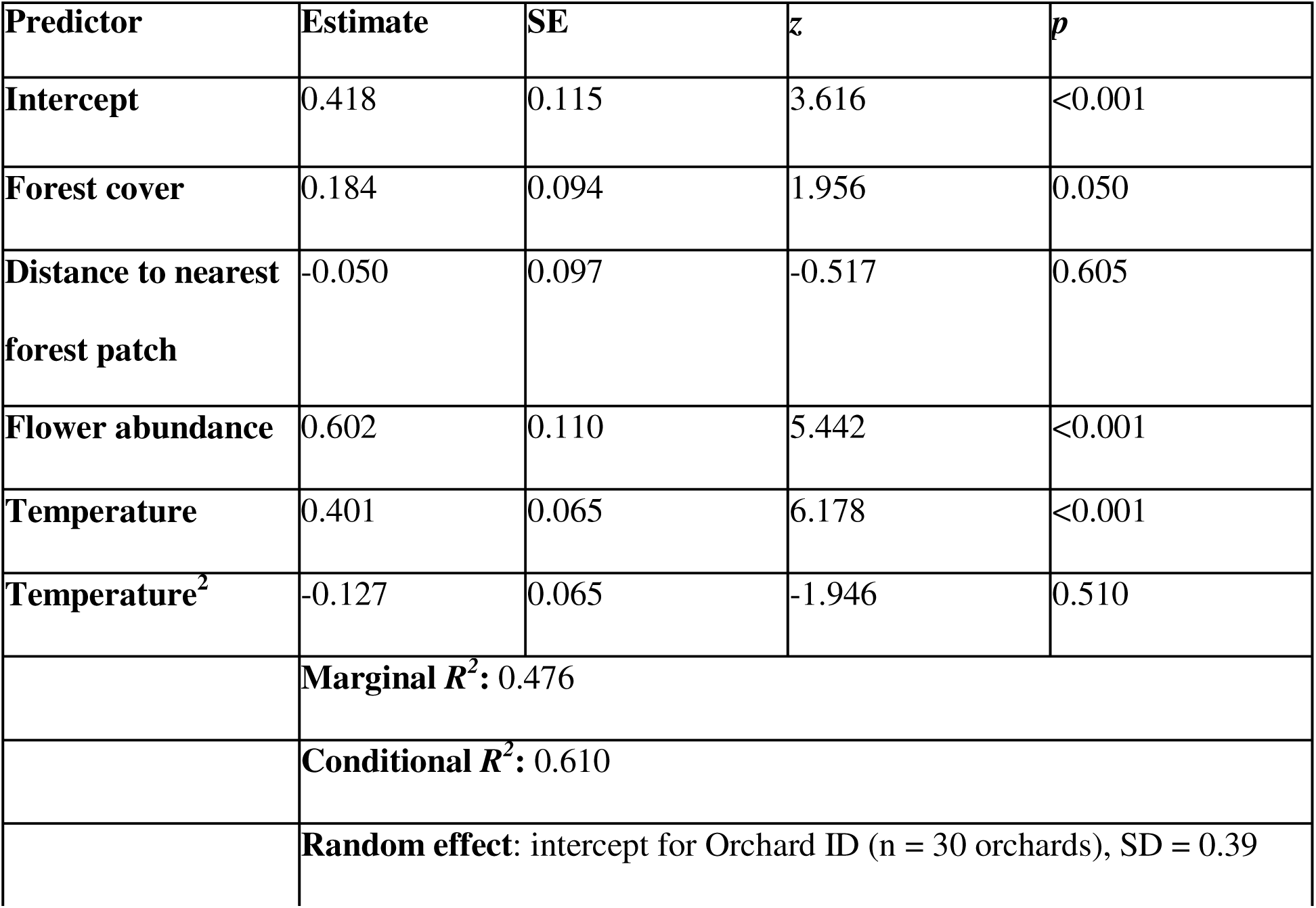
Output of the generalised linear mixed model (negative-binomial error structure) that examined the relationship between pollinator richness in each session and forest cover (500 m buffer), distance to nearest patch, flower abundance, and temperature (with linear and quadratic terms). We defined orchard ID as the random effect. We also report the marginal and conditional *R^2^*.

| Predictor | Estimate | SE | z | p |
| --- | --- | --- | --- | --- |
| Intercept | 0.418 | 0.115 | 3.616 | <0.001 |
| Forest cover | 0.184 | 0.094 | 1.956 | 0.050 |
| Distance to nearest forest patch | -0.050 | 0.097 | -0.517 | 0.605 |
| Flower abundance | 0.602 | 0.110 | 5.442 | <0.001 |
| Temperature | 0.401 | 0.065 | 6.178 | <0.001 |
| Temperature<sup>2</sup> | -0.127 | 0.065 | -1.946 | 0.510 |
| | Marginal $R^2$: 0.476 | | | |
| | Conditional $R^2$: 0.610 | | | |
|  | Random effect: intercept for Orchard ID (n = 30 orchards), SD = 0.39 |  |  |  |

**Table S4.** Output of the Generalised Linear Model (negative-binomial error structure) that examined the relationship between number of pollinators and forest cover, distance to nearest patch, flower abundance, and temperature (with linear and quadratic terms). We also report the marginal and conditional *R^2^*.

| Predictor | Estimate | SE | z | p |
| --- | --- | --- | --- | --- |
| Intercept | 1.074 | 0.129 | 8.295 | <0.001 |
| Forest cover | 0.224 | 0.104 | 2.137 | 0.032 |
| Distance to nearest forest patch | -0.025 | 0.105 | -0.244 | 0.807 |
| Flower abundance | 0.823 | 0.124 | 6.623 | <0.001 |
| Temperature | 0.156 | 0.085 | 1.835 | 0.066 |
| Temperature<sup>2</sup> | -0.224 | 0.082 | -2.733 | 0.006 |
| | Marginal $R^2$: 0.507 | | | |
| | Conditional $R^2$: 0.590 | | | |
|  | Random effect: intercept for Orchard ID (n = 30 orchards), SD = 0.36 |  |  |  |

**Table S5.** Output of the Generalised Linear Model (negative-binomial error structure) that examined the relationship between number of visits to cashew flowers and forest cover (500 m buffer), distance to nearest patch, flower abundance, and temperature (with linear and quadratic terms). We also report the marginal and conditional *R^2^*.

| Predictor | Estimate | SE | z | p |
| --- | --- | --- | --- | --- |
| Intercept | 3.438 | 0.211 | 16.294 | <0.001 |
| Forest cover | 0.531 | 0.177 | 2.990 | 0.002 |
| Distance to nearest forest patch | 0.144 | 0.175 | 0.823 | 0.410 |
| Flower abundance | 1.388 | 0.200 | 6.929 | <0.001 |
| Temperature | 0.111 | 0.148 | 0.753 | 0.451 |
| Temperature<sup>2</sup> | -0.41 | 0.132 | -3.110 | 0.001 |
|  | **Marginal $R^2$: 0.598** |  |  |  |
|  | **Conditional $R^2$: 0.670** |  |  |  |
|  | **Random effect:** Intercept for Orchard ID (n = 30 orchards), SD = 0.53 |  |  |  |

## REFERENCES

Aguirre, A., & Dirzo, R. (2008). Effects of fragmentation on pollinator abundance and fruit set of an abundant understory palm in a Mexican tropical forest. Biological Conservation, 141(2), 375–384. 10.1016/j.biocon.2007.09.014

Araújo, E. D., Costa, M., Chaud-Netto, J., & Fowler, H. G. (2004). Body size and flight distance in stingless bees (Hymenoptera: Meliponini): inference of flight range and possible ecological implications. Brazilian Journal of Biology, 64, 563–568. 10.1590/S1519-69842004000400003

Basu, P., Ngo, H. T., Aizen, M. A., Garibaldi, L. A., Gemmill-Herren, B., Imperatriz-Fonseca, V., Klein, A. M., Potts, S. G., Seymour, C. L., & Vanbergen, A. J. (2024). Pesticide impacts on insect pollinators: Current knowledge and future research challenges. Science of The Total Environment, 954, 176656. 10.1016/j.scitotenv.2024.176656

Bentrup, G., Hopwood, J., Adamson, N. L., Powers, R., & Vaughan, M. (2021). The Role of Temperate Agroforestry Practices in Supporting Pollinators. In R. P. Udawatta & S. Jose (Eds), Agroforestry and Ecosystem Services (pp. 275–304). Springer International Publishing. 10.1007/978-3-030-80060-4_11

Bentrup, G., Hopwood, J., Adamson, N. L., & Vaughan, M. (2019). Temperate Agroforestry Systems and Insect Pollinators: A Review. Forests, 10(11), 981. 10.3390/f10110981

Bhattacharya, A. (2004). Flower visitors and fruitset of Anacardium occidentale. Annales Botanici Fennici, 41(No. 6 (2004)), 385–392.

Biswas, N., Biniwale, S., Sadekar, V., Taneja, Y., Kulbhushansingh, S., Osuri, A., & Naniwadekar. (2025). Securing lowland forests is critical for bird conservation in the Western Ghats Biodiversity Hotspot. Oryx, Accepted.

Biswas, N., Sadekar, V., Biniwale, S., Taneja, Y., Osuri, A. M., Page, N., Suryawanshi, K., & Naniwadekar, R. (2024). Chronic disturbance of moist tropical forests favours deciduous over evergreen tree communities across a climate gradient in the Western Ghats (p. 2024.01.16.575954). bioRxiv. 10.1101/2024.01.16.575954

Brunet, J., & Van Etten, M. (2019). The Response of Floral Traits Associated with Pollinator Attraction to Environmental Changes Expected under Anthropogenic Climate Change in High-Altitude Habitats. International Journal of Plant Sciences. 10.1086/705591

Campbell, A. J., Lichtenberg, E. M., Carvalheiro, L. G., Menezes, C., Borges, R. C., Coelho, B. W. T., Freitas, M. A. B., Giannini, T. C., Leão, K. L., de Oliveira, F. F., Silva, T. S. F., & Maués, M. M. (2022). High bee functional diversity buffers crop pollination services against Amazon deforestation. Agriculture, Ecosystems & Environment, 326, 107777. 10.1016/j.agee.2021.107777

Cant, E. T., Smith, A. D., Reynolds, D. R., & Osborne, J. L. (2005). Tracking butterfly flight paths across the landscape with harmonic radar. Proceedings of the Royal Society B: Biological Sciences. 10.1098/rspb.2004.3002

Celebrezze, T., & Paton, D. C. (2004). Do introduced honeybees ( *Apis mellifera*, Hymenoptera) provide full pollination service to bird adapted Australian plants with small flowers? An experimental study of *Brachyloma ericoides* (Epacridaceae). Austral Ecology, 29(2), 129–136. 10.1111/j.1442-9993.2003.01328.x

Centeno-Alvarado, D., Lopes, A. V., & Arnan, X. (2024). Shaping pollinator diversity through coffee agroforestry management: A meta-analytical approach. Insect Conservation and Diversity, 17(5), 729–742. 10.1111/icad.12755

Chhaya, K., Nulkar, G., Bapat, P., Barve, H., Bhave, N., Bokkasa, A., Dandekar, V., Desai, S., Ghate, K., Gindi, R., Joglekar, A., Joshi, A., Joshi, P., Kamat, S., Karandikar, M., Khod, C., Kulkarni, M., Lad, H., Mandavkar, A.,…Naniwadekar, R. (2025). Understudied and underprotected: Biodiversity and conservation challenges in the Konkan region of Maharashtra. Ecology. 10.1101/2025.05.16.653652

Corlett, R. T. (2004). Flower visitors and pollination in the Oriental (Indomalayan) Region. Biological Reviews, 79(3), 497–532. 10.1017/S1464793103006341

de Frenne, P., Zellweger, F., Rodríguez-Sánchez, F., Scheffers, B., Hylander, K., Luoto, M., Vellend, M., Verheyen, K., & Lenoir, J. R. M. H. (2019). Global buffering of temperatures under forest canopies. Nature Ecology & Evolution, 3(5), 744–749. 10.1038/s41559-019-0842-1

de Manincor, N., Fisogni, A., & Rafferty, N. (2023). Warming of experimental plant–pollinator communities advances phenologies, alters traits, reduces interactions and depresses reproduction. Ecology Letters, 26, 323–334. 10.1111/ele.14158

Descamps, C., Jambrek, A., Quinet, M., & Jacquemart, A.-L. (2021). Warm Temperatures Reduce Flower Attractiveness and Bumblebee Foraging. Insects, 12(6), 493. 10.3390/insects12060493

Dover, J. W., Sparks, T. H., & Greatorex-Davies, J. N. (1997). The importance of shelter for butterflies in open landscapes. Journal of Insect Conservation, 1(2), 89–97. 10.1023/A:1018487127174

Ewers, R. M., & Banks-Leite, C. (2013). Fragmentation Impairs the Microclimate Buffering Effect of Tropical Forests. PLOS ONE, 8(3), e58093. 10.1371/journal.pone.0058093

Freitas, B. M., Filho, A. J. S. P., Andrade, P. B., Lemos, C. Q., Rocha, E. E. M., Pereira, N. O., Bezerra, A. D. M., Nogueira, D. S., Alencar, R. L., Rocha, R. F., & Mendonça, K. S. (2014). Forest remnants enhance wild pollinator visits to cashew flowers and mitigate pollination deficit in NE Brazil. Journal of Pollination Ecology, 12, 22–30. 10.26786/1920-7603(2014)10

Freitas, B., & Paxton, R. (1996). The role of wind and insects in cashew ( Anacardium occidentale) pollination in NE Brazil. The Journal of Agricultural Science, 126, 319–326. 10.1017/S0021859600074876

Freitas, B., Paxton, R., & holanda neto, joão paulo de. (2002). Identifying pollinators among an array of flower visitors, and the case of inadequate cashew pollination in NE Brazil.

Galbraith, S. M., Cane, J. H., Moldenke, A. R., & Rivers, J. W. (2019). Salvage logging reduces wild bee diversity, but not abundance, in severely burned mixed-conifer forest. Forest Ecology and Management, 453, 117622. 10.1016/j.foreco.2019.117622

Ganuza, C., Redlich, S., Uhler, J., Tobisch, C., Rojas-Botero, S., Peters, M. K., Zhang, J., Benjamin, C. S., Englmeier, J., Ewald, J., Fricke, U., Haensel, M., Kollmann, J., Riebl, R., Uphus, L., Müller, J., & Steffan-Dewenter, I. (2022). Interactive effects of climate and land use on pollinator diversity differ among taxa and scales. Science Advances, 8(18), eabm9359. 10.1126/sciadv.abm9359

Garibaldi, L. A., Steffan-Dewenter, I., Kremen, C., Morales, J. M., Bommarco, R., Cunningham, S. A., Carvalheiro, L. G., Chacoff, N. P., Dudenhöffer, J. H., Greenleaf, S. S., Holzschuh, A., Isaacs, R., Krewenka, K., Mandelik, Y., Mayfield, M. M., Morandin, L. A., Potts, S. G., Ricketts, T. H., Szentgyörgyi, H.,…Klein, A. M. (2011). Stability of pollination services decreases with isolation from natural areas despite honey bee visits: Habitat isolation and pollination stability. Ecology Letters, 14(10), 1062–1072. 10.1111/j.1461-0248.2011.01669.x

Garibaldi, L. A., Steffan-Dewenter, I., Winfree, R., Aizen, M. A., Bommarco, R., Cunningham, S. A., Kremen, C., Carvalheiro, L. G., Harder, L. D., Afik, O., Bartomeus, I., Benjamin, F., Boreux, V., Cariveau, D., Chacoff, N. P., Dudenhöffer, J. H., Freitas, B. M., Ghazoul, J., Greenleaf, S.,…Klein, A. M. (2013). Wild Pollinators Enhance Fruit Set of Crops Regardless of Honey Bee Abundance. Science, 339(6127), 1608–1611. 10.1126/science.1230200

Ghazoul, J. (2006). Floral diversity and the facilitation of pollination. Journal of Ecology, 94(2), 295–304. 10.1111/j.1365-2745.2006.01098.x

González Chaves, A., Jaffé, R., Metzger, J., & Kleinert, A. (2020). Forest proximity rather than local forest cover affects bee diversity and coffee pollination services. Landscape Ecology, 35. 10.1007/s10980-020-01061-1

Goodwin, E. K., Rader, R., Encinas-Viso, F., & Saunders, M. E. (2021). Weather Conditions Affect the Visitation Frequency, Richness and Detectability of Insect Flower Visitors in the Australian Alpine Zone. Environmental Entomology, 50(2), 348–358. 10.1093/ee/nvaa180

Hansen, D. M., Olesen, J. M., & Jones, C. G. (2002). Trees, birds and bees in Mauritius: Exploitative competition between introduced honey bees and endemic nectarivorous birds? Journal of Biogeography, 29(5–6), 721–734. 10.1046/j.1365-2699.2002.00720.x

Häussler, J., Sahlin, U., Baey, C., Smith, H., & Clough, Y. (2017). Pollinator population size and pollination ecosystem service responses to enhancing floral and nesting resources. Ecology and Evolution, 7. 10.1002/ece3.2765

Hegland, S., & Boeke, L. (2006). Relationships between the density and diversity of floral resources and flower visitor activity in a temperate grassland community. Ecological Entomology, 31, 532–538. 10.1111/j.1365-2311.2006.00812.x

Hegland, S. J., Nielsen, A., Lázaro, A., Bjerknes, A., & Totland, Ø. (2009). How does climate warming affect plant pollinator interactions? Ecology Letters, 12(2), 184–195. 10.1111/j.1461-0248.2008.01269.x

Hesselbarth, M. H. K., Sciaini, M., Nowosad, J., Hanss, S., structure), L. J. G. (Input on package, structure), J. H. (Input on package, structure), K. A. W. (Input on package, function), F. P. (Original author of underlying C. code for get_nearestneighbour(), lsm_p_circle), P. N. (Original author of underlying C. code for get_circumscribingcircle and, & sample_metrics()), M. S.-M. (Bugfix in. (2025). landscapemetrics: Landscape Metrics for Categorical Map Patterns (Version 2.2.1) [Computer software]. https://cran.r-project.org/web/packages/landscapemetrics/index.html

Hijmans, R. J., Etten, J. van, Sumner, M., Cheng, J., Baston, D., Bevan, A., Bivand, R., Busetto, L., Canty, M., Fasoli, B., Forrest, D., Ghosh, A., Golicher, D., Gray, J., Greenberg, J. A., Hiemstra, P., Hingee, K., Ilich, A., Geosciences, I. for M. A.,…Wueest, R. (2025). raster: Geographic Data Analysis and Modeling (Version 3.6-32) [Computer software]. https://cran.r-project.org/web/packages/raster/index.html

Jevanandam, N., Goh, A. G. R., & Corlett, R. T. (2013). Climate warming and the potential extinction of fig wasps, the obligate pollinators of figs. Biology Letters, 9(3), 20130041. 10.1098/rsbl.2013.0041

Jha, S., & Vandermeer, J. H. (2010). Impacts of coffee agroforestry management on tropical bee communities. Biological Conservation, 143(6), 1423–1431. 10.1016/j.biocon.2010.03.017

Jithin, V., Rane, M., Watve, A., Giri, V. B., & Naniwadekar, R. (2023). Between a rock and a hard place: Comparing rock-dwelling animal prevalence across abandoned paddy, orchards, and rock outcrops in a biodiversity hotspot. Global Ecology and Conservation, 46, e02582. 10.1016/j.gecco.2023.e02582

Jithin, V., Rane, M., Watve, A., & Naniwadekar, R. (2025). Orchards and paddy differentially impact rock outcrop amphibians: Insights from community and species level responses. Ecological Applications, 35(1), e3058. 10.1002/eap.3058

Kalaivanan, D. (2012). Cashew Industry in India – An Overview. CHRONICA HORTICULTURAE, 52, 27–33.

Kalin, M., Armesto, J. J., & Primack, R. (1985). Community studies in pollination ecology in the high temperate Andes of Central Chile. II. Effect of temperature on visitation rates and pollination possibilities. Plant Systematics and Evolution, 149, 187–203. 10.1007/BF00983305

Kapoor, H., & Sharma, S. (2022). Advances in Agricultural and Horticultural Sciences (pp. 591–595).

Kearns, C. A., Inouye, D. W., & Waser, N. M. (1998). ENDANGERED MUTUALISMS: The Conservation of Plant-Pollinator Interactions. Annual Review of Ecology and Systematics, 29(1), 83–112. 10.1146/annurev.ecolsys.29.1.83

Kennedy, C. M., Lonsdorf, E., Neel, M. C., Williams, N. M., Ricketts, T. H., Winfree, R., Bommarco, R., Brittain, C., Burley, A. L., Cariveau, D., Carvalheiro, L. G., Chacoff, N. P., Cunningham, S. A., Danforth, B. N., Dudenhöffer, J., Elle, E., Gaines, H. R., Garibaldi, L. A., Gratton, C.,…Kremen, C. (2013). A global quantitative synthesis of local and landscape effects on wild bee pollinators in agroecosystems. Ecology Letters, 16(5), 584–599. 10.1111/ele.12082

Klein, A., Steffan Dewenter, I., Buchori, D., & Tscharntke, T. (2002). Effects of Land Use Intensity in Tropical Agroforestry Systems on Coffee Flower Visiting and Trap Nesting Bees and Wasps. Conservation Biology, 16(4), 1003–1014. 10.1046/j.1523-1739.2002.00499.x

Klein, A.-M., Vaissière, B. E., Cane, J. H., Steffan-Dewenter, I., Cunningham, S. A., Kremen, C., & Tscharntke, T. (2006). Importance of pollinators in changing landscapes for world crops. Proceedings of the Royal Society B: Biological Sciences, 274(1608), 303–313. 10.1098/rspb.2006.3721

Koetz, A. H. (2013). Ecology, Behaviour and Control of Apis cerana with a Focus on Relevance to the Australian Incursion. Insects, 4(4), 558–592. 10.3390/insects4040558

Lad, H., Gosavi, N., Jithin, V., & Naniwadekar, R. (2025). Effects of land use change and elevation on endemic shrub frogs in a biodiversity hotspot. Animal Conservation, 28(3), 389–400. 10.1111/acv.12991

Li, Y., Zhao, M., Motesharrei, S., Mu, Q., Kalnay, E., & Li, S. (2015). Local cooling and warming effects of forests based on satellite observations. Nature Communications, 6(1), 6603. 10.1038/ncomms7603

Luppi, M., Dondina, O., Orioli, V., & Bani, L. (2018). Local and landscape drivers of butterfly richness and abundance in a human-dominated area. Agriculture, Ecosystems & Environment, 254, 138–148. 10.1016/j.agee.2017.11.020

Madhu, N., Sadekar, V., Biswas, N., Jayapal, R., Rege, A., & Naniwadekar, R. (2025a). Native trees within plantations and surrounding forest cover are essential for bird conservation in cashew-dominated landscapes within a biodiversity hotspot. Forest Ecology and Management, 593, 122878. 10.1016/j.foreco.2025.122878

Madhu, N., Sadekar, V., Biswas, N., Jayapal, R., Rege, A., & Naniwadekar, R. (2025b). Effects of land-use intensification on nectar resources, bird assemblages and bird-flower interaction networks in the Western Ghats, India [Preprint]. bioRxiv.

Magrach, A., & Ghazoul, J. (2015). Climate and Pest-Driven Geographic Shifts in Global Coffee Production: Implications for Forest Cover, Biodiversity and Carbon Storage. PLOS ONE, 10(7), e0133071. 10.1371/journal.pone.0133071

Majewska, A. A., & Altizer, S. (2020). Planting gardens to support insect pollinators. Conservation Biology, 34(1), 15–25. 10.1111/cobi.13271

Makino, T. T., Ohashi, K., & Sakai, S. (2007). How do floral display size and the density of surrounding flowers influence the likelihood of bumble bee revisitation to a plant? Functional Ecology, 21(1), 87–95. 10.1111/j.1365-2435.2006.01211.x

Marini, L., Quaranta, M., Fontana, P., Biesmeijer, J. C., & Bommarco, R. (2012a). Landscape context and elevation affect pollinator communities in intensive apple orchards. Basic and Applied Ecology, 13(8), 681–689. 10.1016/j.baae.2012.09.003

Marini, L., Quaranta, M., Fontana, P., Biesmeijer, J. C., & Bommarco, R. (2012b). Landscape context and elevation affect pollinator communities in intensive apple orchards. Basic and Applied Ecology, 13(8), 681–689. 10.1016/j.baae.2012.09.003

Marshall, L., Perdijk, F., Dendoncker, N., Kunin, W., Roberts, S., & Biesmeijer, J. C. (2020). Bumblebees moving up: Shifts in elevation ranges in the Pyrenees over 115 years. Proceedings of the Royal Society B: Biological Sciences, 287(1938), 20202201. 10.1098/rspb.2020.2201

McDermott Long, O., Warren, R., Price, J., Brereton, T., Botham, M., & Franco, A. (2016). Sensitivity of UK butterflies to local climatic extremes: Which life stages are most at risk? The Journal of Animal Ecology, 86. 10.1111/1365-2656.12594

Millard, J., Outhwaite, C. L., Kinnersley, R., Freeman, R., Gregory, R. D., Adedoja, O., Gavini, S., Kioko, E., Kuhlmann, M., Ollerton, J., Ren, Z.-X., & Newbold, T. (2021). Global effects of land-use intensity on local pollinator biodiversity. Nature Communications, 12(1), 2902. 10.1038/s41467-021-23228-3

Mir, J., Kumar, D., & Pal, A. (2014). Physiology of Flowering in Perennial Temperate Fruit Crops.

Munje, A., & Kumar, A. (2022). Bird community structure in a mixed forest-production landscape in the northern Western Ghats, India. 10.1101/2022.04.04.486917

Nayak, M., & Paled, M. (2018). Trends in Area, Production, Yield and Export-Import of Cashew in India-An Economic Analysis. International Journal of Current Microbiology and Applied Sciences, 7, 1088–1098. 10.20546/ijcmas.2018.712.135

Ohashi, K., & Yahara, T. (2001). Behavioural responses of pollinators to variation in floral display size and their influences on the evolution of floral traits. In L. Chittka & J. D. Thomson (Eds), Cognitive Ecology of Pollination (1st edn, pp. 274–296). Cambridge University Press. 10.1017/CBO9780511542268.015

Ollerton, J., Winfree, R., & Tarrant, S. (2011). How many flowering plants are pollinated by animals? Oikos, 120(3), 321–326. 10.1111/j.1600-0706.2010.18644.x

Ovaskainen, O., & Abrego, N. (2020). Joint Species Distribution Modelling: With Applications in R. Cambridge University Press. 10.1017/9781108591720

Pebesma, E., Bivand, R., Racine, E., Sumner, M., Cook, I., Keitt, T., Lovelace, R., Wickham, H., Ooms, J., Müller, K., Pedersen, T. L., Baston, D., & Dunnington, D. (2025). sf: Simple Features for R (Version 1.0-21) [Computer software]. https://cran.r-project.org/web/packages/sf/index.html

Porto, R. G., De Almeida, R. F., Cruz-Neto, O., Tabarelli, M., Viana, B. F., Peres, C. A., & Lopes, A. V. (2020). Pollination ecosystem services: A comprehensive review of economic values, research funding and policy actions. Food Security, 12(6), 1425–1442. 10.1007/s12571-020-01043-w

Potts, S., Biesmeijer, J. C., Kremen, C., Neumann, P., Schweiger, O., & Kunin, W. E. (2010). Global pollinator declines: Trends, impacts and drivers. Trends in Ecology & Evolution, 25(6), 345–353. 10.1016/j.tree.2010.01.007

Potts, S. G., Biesmeijer, J. C., Kremen, C., Neumann, P., Schweiger, O., & Kunin, W. E. (2010). Global pollinator declines: Trends, impacts and drivers. Trends in Ecology & Evolution, 25(6), 345–353. 10.1016/j.tree.2010.01.007

Potts, S., Petanidou, T., Roberts, S., O’Toole, C., Hulbert, A., & Willmer, P. (2006). Plant-pollinator biodiversity and pollination services in a complex Mediterranean landscape. Biological Conservation, 129(4), 519–529. 10.1016/j.biocon.2005.11.019

Potts, S., Vulliamy, B., Dafni, A., Ne’eman, G., & Willmer, P. (2003). Linking bees and flowers: How do floral communities structure pollinator communities? Ecology, 84, 2628–2642. 10.1890/02-0136

Proesmans, W., Bonte, D., Smagghe, G., Meeus, I., Decocq, G., Spicher, F., Kolb, A., Lemke, I., Diekmann, M., Bruun, H., Wulf, M., Van Den Berge, S., & Verheyen, K. (2019). Small forest patches as pollinator habitat: Oases in an agricultural desert? Landscape Ecology, 34. 10.1007/s10980-019-00782-2

Punchihewa, R. W. K., Koeniger, N., Kevan, P. G., & Gadawski, R. M. (1985). Observations on the Dance Communication and Natural Foraging Ranges of Apis Cerana, Apis Dorsata and Apis Florea in Sri Lanka. Journal of Apicultural Research, 24(3), 168–175. 10.1080/00218839.1985.11100667

Pyke, G. H., Thomson, J. D., Inouye, D. W., & Miller, T. J. (2016). Effects of climate change on phenologies and distributions of bumble bees and the plants they visit. Ecosphere, 7(3), e01267. 10.1002/ecs2.1267

Rafferty, N. E. (2017). Effects of global change on insect pollinators: Multiple drivers lead to novel communities. Current Opinion in Insect Science, 23, 22–27. 10.1016/j.cois.2017.06.009

Rege, A., Bodhankar Warnekar, S., & Lee, J. S. H. (2022). Mapping cashew monocultures in the Western Ghats using optical and radar imagery in Google Earth Engine. Remote Sensing Applications: Society and Environment, 28, 100861. 10.1016/j.rsase.2022.100861

Rege, A., & Lee, J. S. H. (2022). State-led agricultural subsidies drive monoculture cultivar cashew expansion in northern Western Ghats, India. PLOS ONE, 17(6), e0269092. 10.1371/journal.pone.0269092

Rege, A., Warnekar, S. B., & Lee, J. S. H. (2022). Mapping cashew monocultures in the Western Ghats using optical and radar imagery in Google Earth Engine. Remote Sensing Applications: Society and Environment, 28, 100861. 10.1016/j.rsase.2022.100861

Sadekar, V., Satish, A., & Naniwadekar, R. (2025). Cashew orchard soil properties, Dodamarg, Northern Western Ghats, India [Data set]. Zenodo. 10.5281/zenodo.16017208

Scaven, V. L., & Rafferty, N. E. (2013). Physiological effects of climate warming on flowering plants and insect pollinators and potential consequences for their interactions. Current Zoology, 59(3), 418–426. 10.1093/czoolo/59.3.418

Seeley, T. D., Seeley, R. H., & Akratanakul, P. (1982). Colony Defense Strategies of the Honeybees in Thailand. Ecological Monographs, 52(1), 43–63. 10.2307/2937344

Simanonok, M., Powley, M., & Otto, C. (2022). Cavity-nesting Bee Nesting Success Across Gradients of Floral Resources and Land Cover.

Smith, T. J., & Mayfield, M. M. (2018). The effect of habitat fragmentation on the bee visitor assemblages of three Australian tropical rainforest tree species. Ecology and Evolution, 8(16), 8204–8216. 10.1002/ece3.4339

Staton, T., Walters, R. J., Smith, J., & Girling, R. D. (2019). Evaluating the effects of integrating trees into temperate arable systems on pest control and pollination. Agricultural Systems, 176, 102676. 10.1016/j.agsy.2019.102676

Steffan-Dewenter, I., Munzenberg, U., Burger, C., Thies, C., & Tscharntke, T. (2002). Scale-Dependent Effects of Landscape Context on Three Pollinator Guilds. Ecology, 83, 1421–1432. 10.2307/3071954

Steffan-Dewenter, I., Potts, S. G., & Packer, L. (2005). Pollinator diversity and crop pollination services are at risk. Trends in Ecology & Evolution, 20(12), 651–652. 10.1016/j.tree.2005.09.004

Tikhonov, G., Ovaskainen, O., Oksanen, J., Jonge, M. de, Opedal, O., & Dallas, T. (2025). Hmsc: Hierarchical Model of Species Communities (Version 3.3-7) [Computer software]. https://cran.r-project.org/web/packages/Hmsc/index.html

Uhl, P., & Brühl, C. A. (2019). The Impact of Pesticides on Flower Visiting Insects: A Review with Regard to European Risk Assessment. Environmental Toxicology and Chemistry, 38(11), 2355–2370. 10.1002/etc.4572

Ulyshen, M., Urban-Mead, K., Dorey, J., & Rivers, J. (2023). Forests are critically important to global pollinator diversity and enhance pollination in adjacent crops. Biological Reviews of the Cambridge Philosophical Society, 98. 10.1111/brv.12947

Vanitha, K., & Raviprasad, T. N. (2019). Diversity, Species Richness and Foraging Behaviour of Pollinators in Cashew. Agricultural Research, 8(2), 197–206. 10.1007/s40003-018-0370-2

Vaughton, G. (1996). Pollination disruption by European honeybees in the Australian bird-pollinated shrubGrevillea barklyana (Proteaceae). Plant Systematics and Evolution, 200(1–2), 89–100. 10.1007/BF00984750

Vázquez, D. P., Alvarez, J. A., Debandi, G., Aranibar, J. N., & Villagra, P. E. (2011). Ecological consequences of dead wood extraction in an arid ecosystem. Basic and Applied Ecology, 12(8), 722–732. 10.1016/j.baae.2011.08.009

Vicens, N., & Bosch, J. (2000). Weather-Dependent Pollinator Activity in an Apple Orchard, with Special Reference to Osmia cornuta and Apis mellifera (Hymenoptera: Megachilidae and Apidae). Environmental Entomology, 29, 413–420. 10.1603/0046-225X-29.3.413

Vizentin-Bugoni, J., Maruyama, P. K., de Souza, C. S., Ollerton, J., Rech, A. R., & Sazima, M. (2018). Plant-Pollinator Networks in the Tropics: A Review. In W. Dáttilo & V. Rico-Gray (Eds), Ecological Networks in the Tropics: An Integrative Overview of Species Interactions from Some of the Most Species-Rich Habitats on Earth (pp. 73–91). Springer International Publishing. 10.1007/978-3-319-68228-0_6

Von Arx, G., Graf Pannatier, E., Thimonier, A., & Rebetez, M. (2013). Microclimate in forests with varying leaf area index and soil moisture: Potential implications for seedling establishment in a changing climate. Journal of Ecology, 101(5), 1201–1213. 10.1111/1365-2745.12121

Wagner, D. L. (2020). Insect Declines in the Anthropocene. Annual Review of Entomology, 65(Volume 65, 2020), 457–480. 10.1146/annurev-ento-011019-025151

Whittaker, R. H. (1972). EVOLUTION AND MEASUREMENT OF SPECIES DIVERSITY. TAXON, 21(2–3), 213–251. 10.2307/1218190

Williams, P. H., Araújo, M. B., & Rasmont, P. (2007). Can vulnerability among British bumblebee (Bombus) species be explained by niche position and breadth? Biological Conservation, 138(3–4), 493–505. 10.1016/j.biocon.2007.06.001

Willmer, P. G. (1983). Thermal constraints on activity patterns in nectar feeding insects. Ecological Entomology, 8(4), 455–469. 10.1111/j.1365-2311.1983.tb00524.x

Willmer, P. G., & Stone, G. N. (2004). Behavioral, Ecological, and Physiological Determinants of the Activity Patterns of Bees. In Advances in the Study of Behavior (Vol. 34, pp. 347–466). Elsevier. 10.1016/S0065-3454(04)34009-X

Zurbuchen, A., Landert, L., Klaiber, J., Müller, A., Hein, S., & Dorn, S. (2010). Maximum foraging ranges in solitary bees: Only few individuals have the capability to cover long foraging distances. Biological Conservation, 143(3), 669–676. 10.1016/j.biocon.2009.12.003

